# Alphafold-multimer predicts cross-kingdom interactions at the plant-pathogen interface

**DOI:** 10.1101/2023.04.03.535425

**Authors:** Felix Homma, Jie Huang, Renier A. L. van der Hoorn

## Abstract

Adapted plant pathogens from various microbial kingdoms produce hundreds of unrelated small secreted proteins (SSPs) with elusive roles. Some of these SSPs might be inhibitors targeting the most harmful hydrolases secreted by the host. Here, we used Alphafold-Multimer (AFM) to screen 1,879 SSPs of seven tomato pathogens for interacting with six defence-related hydrolases of tomato that accumulate to high levels in the apoplast during infection. This screen of 11,274 protein pairs identified 15 SSPs that are predicted to obstruct the active site of chitinases and proteases with an intrinsic fold. Four SSPs were experimentally verified to be inhibitors of pathogenesis-related subtilase P69B, including extracellular protein-36 (Ecp36) and secreted-into-xylem-15 (Six15) of the fungal tomato pathogens *Cladosporium fulvum* and *Fusarium oxysporum*, respectively. Together with a novel P69B inhibitor from the bacterial pathogen *Xanthomonas perforans* and the previously reported Kazal-like inhibitors of the oomycete pathogen *Phytophthora infestans*, P69B emerges as an important effector hub targeted by different microbial kingdoms, consistent with the presence of a hyper-variant residue in P69B orthologs and gene duplication and diversification of P69B paralogs that could avoid inhibitor binding. This study demonstrates the power of artificial intelligence to accurately predict novel cross-kingdom interactions at the plant-pathogen interface.

## INTRODUCTION

The extracellular space inside plant tissues (the apoplast) is heavily defended (Doehlemann & Hemetsberger, 2013; Darino et al., 2022). In response to apoplast colonization by bacterial, fungal and oomycete pathogens, the host plant secretes a broad diversity of metabolites and proteins that are presumably toxic and harmful to extracellular microbes. Adapted pathogens, however, have learned to live in this challenging environment, but the mechanisms that these pathogens use to avoid or suppress extracellular immunity are largely unknown.

Hydrolytic enzymes, such as proteases, glycosidases and lipases, are abundantly secreted proteins during the plant defense response. Many of these defense-induced hydrolases are described since the 1980s as pathogenesis-related (PR) proteins, as they accumulate to high levels in the apoplast of infected plants (Van Loon et al., 2006). These PR proteins include glucanases (PR2), chitinases (PR3), and proteases (PR7). The PR7 proteases are also called P69 subtilases as they are subtilisin-like proteases that accumulate at ~70 kDa in tomato upon infection with various pathogens (Jorda et al., 1999; Jorda et al., 2000).

The relevance of P69s and other secreted defence-related hydrolases is underlined by the fact that pathogens suppress their activity with pathogen-secreted inhibitors. Tomato P69B subtilase, for instance, is targeted by Kazal-like inhibitors Epi1 and Epi10 from *P. infestans* (Tian et al., 2004; Tian et al., 2005) and the defence-related papain-like Phytophthora-inhibited protease-1 (Pip1) from tomato, is targeted by cystatin-like EpiC1 and EpiC2B of *P. infestans* (Tian et al., 2007). Pip1 is also targeted by Avr2 from the fungal tomato pathogen *Cladosporium fulvum* (syn. *Passalora fulva*; Shabab et al., 2008; Van Esse et al., 2008), and by the chagasin-like Cip1 from the bacterial tomato pathogen *Pseudomonas syringae* pv. *syringae* (Shindo et al., 2016). In all these examples, pathogen-derived inhibitors are small secreted proteins (SSPs) that are often stabilized by disulfide bridges. Additional pathogen-produced SSP targeting host hydrolases include Pit2 from the fungal maize pathogen *Ustilago maydis* (Mueller et al., 2013); and SDE1 from the bacterial citrus pathogen *Liberibacter asiaticus* (Clark et al., 2018).

The targeting of secreted hydrolases by multiple pathogen-produced SSPs implicates that these secreted hydrolases can play important roles in immunity and that adapted pathogens are all secreting inhibitors targeting the most harmful hydrolases. Indeed, Pip1 depletion by RNAi makes tomato hypersusceptible to bacterial, fungal and oomycete pathogens (Ilyas et al., 2015), illustrating that Pip1 provides broad range immunity, despite being targeted by pathogen-derived inhibitors. Following the same narrative, we discovered that plant-secreted beta-galactosidase BGAL1 triggers the release of immunogenic flagellin fragments, a study that was sparked by the discovery that BGAL1 is suppressed during *P. syringae* infection (Buscaill et al., 2019). We have uncovered an additional 59 apoplastic hydrolases that are suppressed during *P. syringae* infection, one of which is *Nb*PR3, a neo-functionalised chitinase that provides antibacterial immunity (Sueldo et al., 2023).

The plant-pathogen arms race between inhibitors and their target hydrolases results in the selection of residues at the interaction interface, as a ‘ring-of-fire’, indicative of a footprint of an arms race with pathogen-derived inhibitors. Examples include Class-I chitinases (Bishop et al., 2000), soybean endoglucanase EGase (Bishop et al., 2005), and tomato papain-like protease Rcr3 (Shabab et el., 2008). Variant residues in Rcr3 indeed interfere with Avr2 binding (Shabab et al., 2008; Kourelis et al., 2020), and variant residues in soybean EGaseA are predicted to interact with variant residues in GIP1 from *Phytophthora sojae* (Damasenco et al., 2008). These discoveries imply that engineering of inhibitor-insensitive hydrolases is feasible and can provide a new crop protection strategy.

New approaches are needed to discover and exploit antagonistic interactions at the plant-pathogen interface. Here, we tested the use of Alphafold Multimer (AFM, Evans et al., 2021) to discover novel extracellular inhibitor-hydrolase interactions. Alphafold2 accurately predicts structures of novel proteins using artificial intelligence trained on multiple sequence alignments (MSA) and structural information (Jumper et al., 2021). Alphafold2 produces a predicted Template Modeling (pTM) score and visualizes the confidence in predicted structures using the predicted local Distance Difference Test (plDDT). AFM is an extension of Alphafold2 developed by DeepMind to predict structures of protein complexes and produces the interface pTM score (ipTM), that weighs heavily in the overall score of predicted complexes (0.8ipTM+0.2pTM, Evans et al., 2021). AFM has been used for a variety of predictions, e.g. to confirm and predict protein-protein complexes in yeast (Humphreys et al., 2021); or to predict typical and atypical ATG8 binding motifs in eukaryote proteins (Ibrahim et al., 2023). Here we demonstrate that AFM can also be used for cross-kingdom discovery screens for novel protein-protein interactions at the plant pathogen interface, illustrated with the discovery of four novel pathogen-derived inhibitors targeting a tomato-secreted immune protease P69B.

## RESULTS

### AFM scores distinguish existing from non-existing complexes

To test the prediction of protein complexes at the plant-pathogen interface with AFM, we first predicted two well-studied protein complexes from the interactions between tomato and the late blight pathogen *P. infestans*. The first complex is between the P69B subtilase of domesticated tomato (*Solanum lycopersicum, Sl*) and the first Kazal domain of Epi1 of *P. infestans* (Epi1a) (Tian et al., 2004). Both P69B and Epi1a have high mean non-gap MSA depth (**Figure 1A**) and the best ipTM+pTM score that AFM predicts for P69B-Epi1a is 0.93, supported with high plDDT scores at the interaction interface (**Figure 1B**). The predicted complex is accurate because the Reactive Site Loop (RLS) of Epi1a in the predicted model forms eleven hydrogen bonds in the active site, and the P1=Asp residue of Epi1a nicely fits in the S1 substrate binding pocket of P69B, consistent with how Kazal-like inhibitors bind to subtilases (Lu et al., 2001).

**Figure 1.**
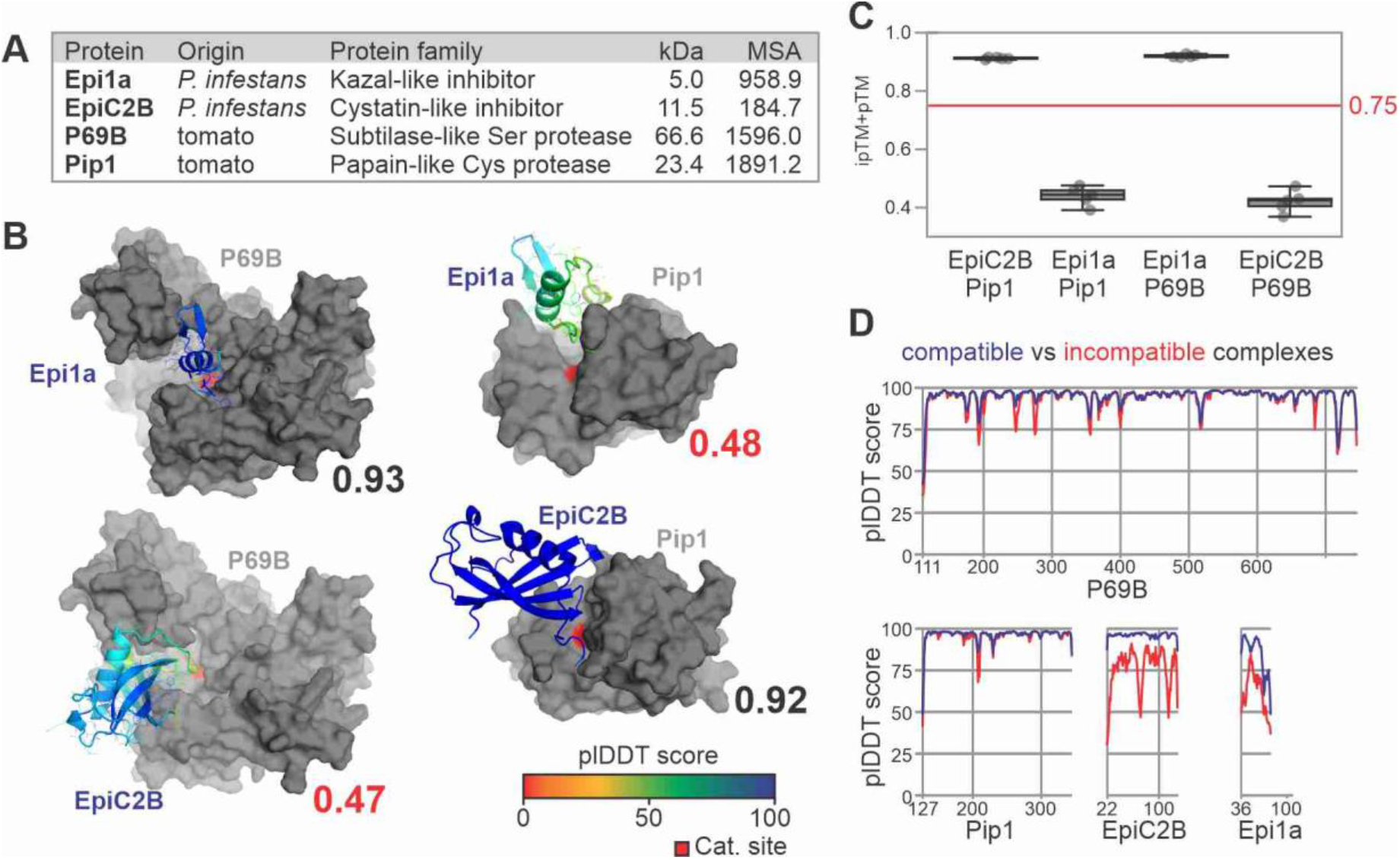
AFM accurately distinguishes existing from non-existing complexes at the plant-pathogen interface. **(A)** Used proteins with their nature and origin, mature molecular weight (MW, in kDa) and depth of multiple sequence alignment (MSA) detected for proteins in compatible complexes. **(B)** Best structures predicted by AFM for existing and non-existing inhibitor-hydrolase complexes, with their iTM+ipTM scores ranging from 0 (worst) to 1 (best). Pip1 and P69B are shown in grey, with their catalytically active residue in red. EpiC2B and Epi1a are coloured using a rainbow scheme based on their plDDT scores, which range from 0 (worst) to 100 (best). **(C)** ipTM+pTM quality scores for each of the five models for each of the protein pairs, showing the median, 25th and 75th percentiles, and whiskers representing 1.5 times the interquartile range. **(D)** plDDT scores within the four proteins in predicted compatible (blue) and incompatible (red) complexes.

The second complex is between the papain-like protease Pip1 of tomato and the cystatin-like EpiC2B of *P. infestans* (Tian et al., 2007). Also these two proteins have high mean non-gap MSA depth (**Figure 1A**), and the best AFM-predicted model has a high combined ipTM+pTM score of 0.92, supported by high plDDT scores at the interaction interface (**Figure 1B**). As expected, the tripartite wedge of EpiC2B occupies the substrate binding groove of Pip1 and forms 13 predicted hydrogen bonds with Pip1, consistent with the literature on cystatin-papain interactions (Chu et al., 2011).

Taking advantage of the fact that P69B and Pip1 are unrelated proteases, and Epi1a and EpiC2B are unrelated inhibitors, we next tested if AFM would produce different scores with incompatible protein pairs by swapping the inhibitors between the proteases. Indeed, the best ipTM+pTM scores are now much lower for these incompatible complexes: 0.47 for P69B-EpiC2B and 0.47 for Pip1-Epi1a, respectively. The individual proteins are still folded as expected, and occupy the substrate binding grooves (**Figure 1B**). The plDTT scores were reduced in incompatible complexes for whole inhibitors, and for multiple sites in the proteases (**Figure 1D**). For each of the four protein pairs, all five AFM-predicted models were consistently assigned similar ipTM+pTM (**Figure 1C**), facilitating statistical analysis that demonstrates that AFM scores are statistically different between compatible and incompatible complexes (p= 2.1e-09 and 1.8e-9, for P69B and Pip1, respectively).

### AFM screen between 1,879 SSPs and 6 hydrolases identifies 376 candidate complexes

Having established that AFM is able to distinguish between compatible and incompatible complexes, we decided to use AFM as an interactomic discovery platform to identify novel pathogen-derived inhibitors targeting extracellular defence-related hydrolases of tomato, based on the hypothesis that all tomato pathogens will secrete inhibitors targeting harmful hydrolases. We selected 1,879 small secreted proteins (SSPs) from seven different tomato pathogens representing different kingdoms (**Figure 2A**). We included three bacterial tomato pathogens: *Pseudomonas syringae (Ps), Xanthomonas perforans (Xp*), and *Ralstonia solanacearum (Rs);* three fungal tomato pathogens: *Botrytis cinerea* (*Bc*), *Fusarium oxysporum* f. sp. *lycopersici* (*Fo*), and *Cladosporium fulvum* (*Cf*) and the oomycete pathogen *Phytophthora infestans (Pi). Ps, Xp* and *Cf* are biotrophic leaf pathogens that are exposed to tomato-secreted hydrolases during colonization of the apoplast. *Bc* and *Pi* are hemibiotrophic leaf pathogens that colonize the tomato apoplast during the initial phase of infection. *Rs* and *Fo* colonize the xylem, which is considered part of the apoplast and has a similar content as the leaf apoplast (Houterman et al., 2007). These very different pathogens cause important diseases on tomato (Dean et al. 2012; Mansfield et al. 2012; Kamoun et al. 2015) and their assembled genomes are publicly available (Buell et al., 2003 (*Ps*); Salanoubat et al., 2002 (*Rs*); Van Kan et al., 2017 (*Bc*); Ma et al., 2010 (*Fo);* Zaccaron et al., 2022 (*Cf);* Haas et al., 2009 (*Pi*)). We selected SSPs from these genomes by selecting small proteins (<35 kDa) that have a likely apoplastic localization predicted by either SignalP5.01 or TargetP2.0, supported by ApoplastP1.01 (Sperschneider et al., 2018; Almagro Armenteros et al., 2019a; Almagro Armenteros et al., 2019b). This selection will not include all possible secreted pathogen-derived hydrolase inhibitors, but this number and limited protein size will restrict the AFM screen to a computationally feasible level.

**Figure 2.**
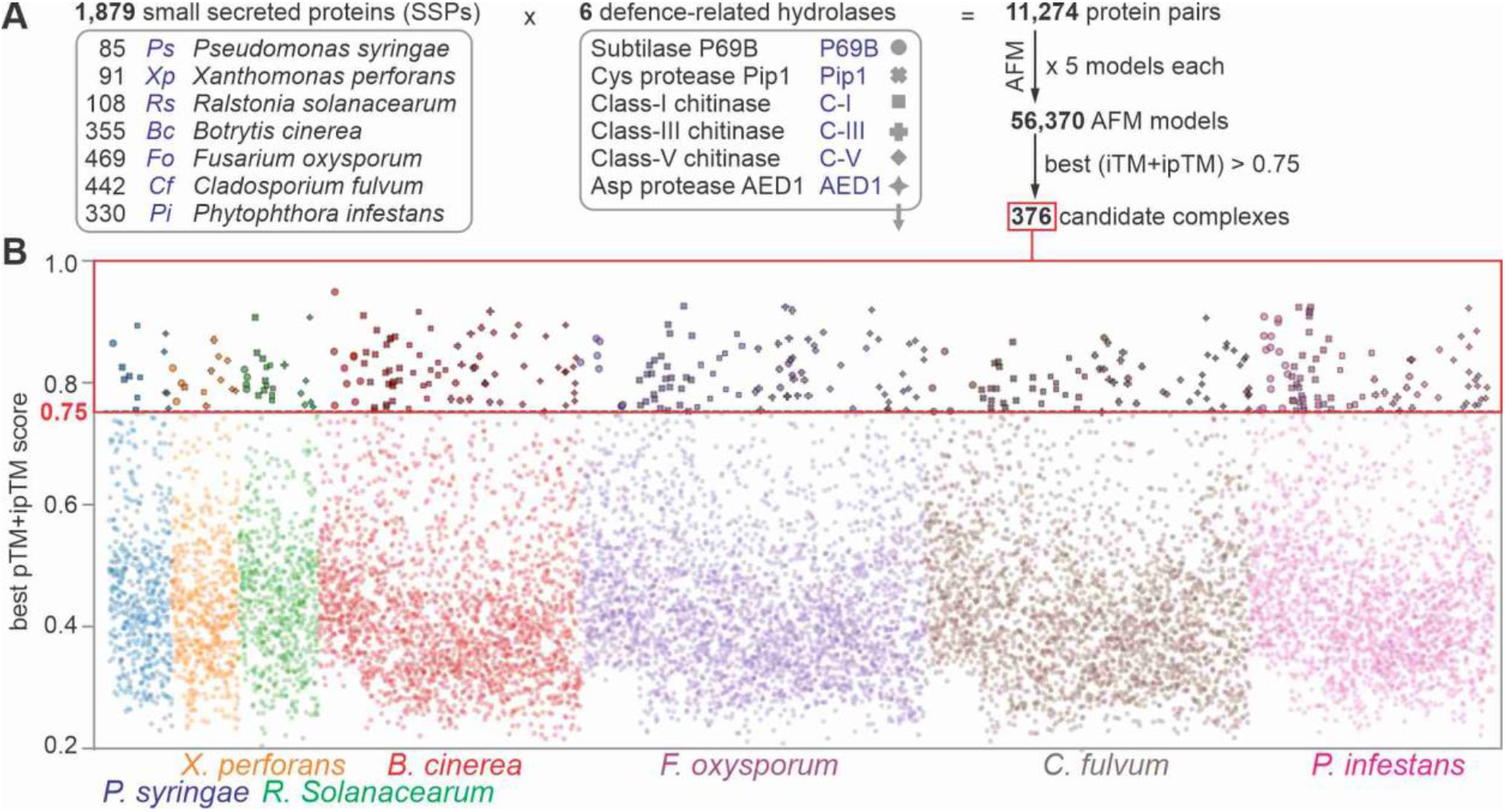
AFM screen for complexes between 1,879 small secreted proteins and 6 hydrolases generated 376 candidate complexes. **(A)** 1,879 proteins from seven tomato pathogens that are likely secreted and small (<35 kDa) were screened for complexes with six secreted defence-related hydrolases of tomato using alphafold2 multimer (AFM). The best of the five generated AFM models for each of the 11,274 protein pairs were selected if the iTM+ipTM score was 0.75 or higher, resulting in 376 putative complexes. **(B)** best pTM+ipTM scores for all the 11,274 complexes involving 1,879 small secreted proteins (SSPs) of the seven tomato pathogens listed on the bottom. Symbols for complexes with the six different hydrolases (explained in (A)) highlight the 376 candidate complexes with pTM+ipTM ≥ 0.75.

We restricted our AFM screen to identify inhibitors of six defence-related extracellular hydrolases of tomato that have a clear substrate binding groove that will aid the selection of hydrolase inhibitors (**Figure 2A**). Besides P69B and Pip1, we included defence-induced chitinases of classes I, III and V. These are abundant and well-described pathogenesis-related PR3 and PR8 proteins accumulating in the apoplast of tomato upon infection (Joosten et al., 1989). We also included a pepsin-like apoplastic enhanced disease susceptibility (AED1) as a negative control as it is implicated in the suppression of systemic acquired resistance and therefore unlikely to be targeted by inhibitors (SAR, Breitenbach et al., 2014).

We next tested 11,274 protein pairs between the 1,879 SSPs and the six hydrolases using a custom-made AFM workflow where we reduced computing time by avoiding redundant database searches for the same protein. The AFM screen required 13,244 CPU hours (1.51 CPU years) and 8,118 GPU hours (0.93 GPU years), which equals to 1.17 CPU hours and 0.72 GPU hours per protein pair. These hardware requirements were made feasible using the Advanced Research Computing (ARC) facility of the University of Oxford.

The AFM screen resulted in 376 protein pairs with a best ipTM+pTM score of ≥ 0.75 (**Figure 2A**). These 376 protein pairs represent 3.3% of the tested protein pairs. This percentage is intuitively high because we expect that most pathogens produce only one or two inhibitors for each hydrolase (41-84 inhibitors in total) but this total number is sufficiently low to investigate individually. The 376 hits were distributed over the pathogens and hydrolases, such that most pathogens had several candidate inhibitors for each hydrolase (**Figure 2B**).

### Further selection of candidates identifies 15 novel putative complexes during infection

To analyse the structures of the best models for each of these 376 protein pairs, we developed a custom script in python to present the surface of the hydrolase structure in grey, with the active site in red and the putative inhibitor as cartoon and lines, coloured using a rainbow scheme based on the plDDT scores. This presentation facilitated a quick classification of how the SSP binds to the hydrolase.

The 376 complexes were classified into four different groups (**Figure 3A**). One group (19 complexes) were nonsense models, where the two polypeptide strands are entangled into each other, which is unlikely when proteins are folded and secreted by different organisms. A second group (137 complexes) has the substrate binding groove fully exposed and the SSP binding elsewhere on the hydrolase. Although some of these SPPs might be allosteric hydrolase regulators, these complexes were not considered further. In the third group (184 complexes), the active site was blocked by the SSP, but the region blocking the active site had no intrinsic structure, but was rather an unstructured strand bound to the substrate binding groove. Some of these SSPs might be substrates when bound to proteases, but these were not considered further. The fourth group (36 complexes) contains structures where the SSP blocks the active site with an intrinsic structure, often involving multiple disulphide bridges and secondary structures. This type of interaction is common for described inhibitor-hydrolase complexes and these complexes were therefore further analysed.

**Figure 3.**
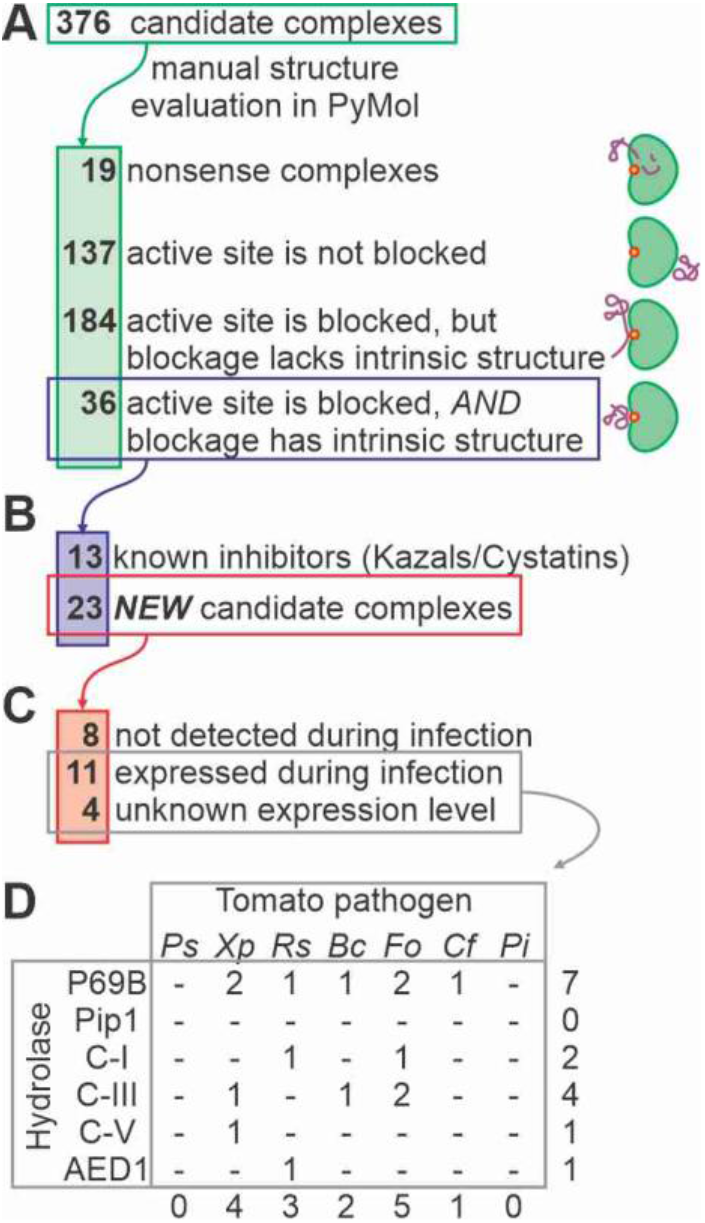
Selection of candidate complexes. **(A)** The 376 candidate complexes were manually screened for complexes where the SSP blocks the active site of the hydrolase with an intrinsic fold. **(B)** The remaining 36 candidate complexes included 13 complexes between Kazal-like inhibitors and P69B and Cystatin-like inhibitors with Pip1, and 23 candidate complexes with other candidate inhibitors. **(C)** Of the 23 remaining candidate complexes, transcriptome analysis of infected plants showed that genes encoding 8 SSPs are not expressed at the tested conditions, whereas genes encoding 11 SSPs are expressed during infection. No transcriptomic datasets were available for *Xp* (4 SSPs). **(D)** Distribution of the final 15 candidate hydrolase-inhibitor complexes over the tested tomato pathogens and target hydrolases.

The 36 complexes included eight complexes of Kazal-like proteins from *Pi* bound to P69B, and five complexes of cystatin-like proteins from *Pi* bound to Pip1 (**Figure 3B**). The selection of these inhibitors validated our manual screening method. However, since these interactions could also be predicted by sequence homology, these were not studied further.

To focus further studies on novel protein complexes that could exist during infection, we mined transcriptomic databases for the expression levels of the remaining 15 inhibitor proteins during infection (Khokhani et al., 2017 (*Rs);* Ilyas et al., 2015 (*Cf);* Müller et al., 2018 (*Bc);* Zhao et al., 2018 (*Fo*)). As most of these studies did not report on pathogen gene expression, we reanalyzed the RNA-seq data by removing plant sequences and mapping the remaining reads against predicted coding sequences of the pathogens, resulting in expression levels for every pathogen in transcript per million (tpm). This way, we identified expression during infection for 11 putative inhibitors, with expression levels ranging from 2.4 to 717 tpm reads (**Figure 3C** and **Table 1**). No transcripts were detected for eight candidate inhibitors. Although the expression of these candidates might have been missed by chosen conditions and materials, these eight candidates were not analyzed further. There was no expression data available for *Xp* infections, but these four candidates were all included.

**Table 1.** 15 novel candidate hydrolase-inhibitor complexes at the plant-pathogen interface

|  |  |  |  | ipTM+pTM |  |  |  |
| --- | --- | --- | --- | --- | --- | --- | --- |
| Sp | Effector<br>accession | Target<br>hydrolase | Annotation | All models | Best<br>model | MSA<br>depth | Expression<br>(tpm) |
| *Xp* | WP008576433.1 | P69B | - | 0.65 ± 0.06 | 0.77 | 96.22 | NA |
| *Xp* | WP046932418.1 | P69B | **P3** *Xp*Ssp1 | 0.64 ± 0.16 | 0.82 | 213.55 | NA |
| *Cf* | KAH3648627.1 | P69B | **P5** *Cf*Ecp36 | 0.55 ± 0.11 | 0.75 | 13.43 | **479 ± 179** |
| *Bc* | XP001545484.1 | P69B | **P6** | 0.57 ± 0.16 | 0.82 | 238.30 | 2.4 ± 0.1 |
| *Rs* | WP011000405.1 | P69B | **P7** | 0.59 ± 0.12 | 0.81 | 614.55 | 6.8 ± 5.7 |
| *Fo* | XP018243121.1 | P69B | **P8** *Fo*TIL | 0.79 ± 0.05 | 0.87 | 815.69 | **683 ± 370** |
| *Fo* | APP91304.1 | P69B | **P9** *Fo*Six15 | 0.69 ± 0.12 | 0.83 | 3.88 | **717 ± 124** |
| *Rs* | WP011001815.1 | C-I | - | 0.50 ± 0.19 | 0.83 | 1798.12 | **599 ± 53** |
| *Fo* | XP018236493.1 | C-I | - | 0.85 ± 0.03 | 0.88 | 355.35 | **216 ± 11** |
| *Xp* | WP046931881.1 | C-III | - | 0.76 ± 0.20 | 0.87 | 141.47 | NA |
| *Bc* | XP001560184.1 | C-III | - | 0.63 ± 0.24 | 0.88 | 296.23 | 59 ± 22 |
| *Fo* | XP018248187.1 | C-III | - | 0.55 ± 0.28 | 0.92 | 171.59 | 5.6 ± 0.2 |
| *Fo* | XP018241286.1 | C-III | - | 0.43 ± 0.23 | 0.87 | 62.22 | 3.9 ± 2.1 |
| *Xp* | WP008572913.1 | C-V | - | 0.49 ± 0.18 | 0.83 | 107.08 | NA |
| *Rs* | WP011002292.1 | AED1 | - | 0.67 ± 0.10 | 0.76 | 576.77 | 2.9 ± 0.2 |

NA, not available

The selection for likely inhibitors that are novel and expressed during infection resulted in 15 proteins that are not equally distributed over the hydrolases and pathogens (**Figure 3D** and **Table 1**). P69B emerges as a putative ‘effector hub’ by being targeted by seven novel putative inhibitors produced by five additional pathogens, in addition to the previously identified Kazal-like inhibitors of *Pi*. No novel inhibitors were identified from pathogens *Ps* and *Pi*, or targeting Pip1. Unexpectedly, putative chitinase inhibitors are also produced by bacterial pathogens, and one putative inhibitor might target AED1.

### Four novel P69B inhibitors were identified by activity labelling

We decided to confirm novel inhibitors of P69B because this hydrolase is targeted by most putative inhibitors and we also have assays available to monitor P69B inhibition. A C-terminally His-tagged P69B was efficiently produced by agroinfiltration of *N. benthamiana* and purified on immobilized Ni-NTA (Paulus et al., 2020). Active-site labelling with fluorescent fluorophosphonate probe FP-TAMRA (Liu et al., 1999) is a sensitive and specific assay to detect P69B inhibition and has been used to confirm that Epi1 inhibits P69B (Paulus et al., 2020).

Six candidate P69B inhibitors were expressed in *E. coli* Rosetta-gami B cells to facilitate the folding of proteins having disulfide bridges. The putative inhibitors were fused to an *N*-terminal double purification tag consisting of a His tag, maltose binding protein (MBP) and a cleavage site for TEV protease (Supplemental **Figure S1**). Two inhibitor candidates did not express sufficiently to pursue the purification. The remaining four fusion proteins were purified over Ni-NTA and amylose resin, subsequently. Next, the purification tag was removed with the TEV protease and the protease and purification tags were removed using the Ni-NTA matrix and 30 kDa centrifugal concentrator, resulting in four purified inhibitor proteins containing only an additional *N*-terminal Gly-Glu-Phe tripeptide (**Figure 4A**). Epi1 (positive control) and EpiC1 (negative control) were produced and purified following the same procedure.

**Figure 4.**
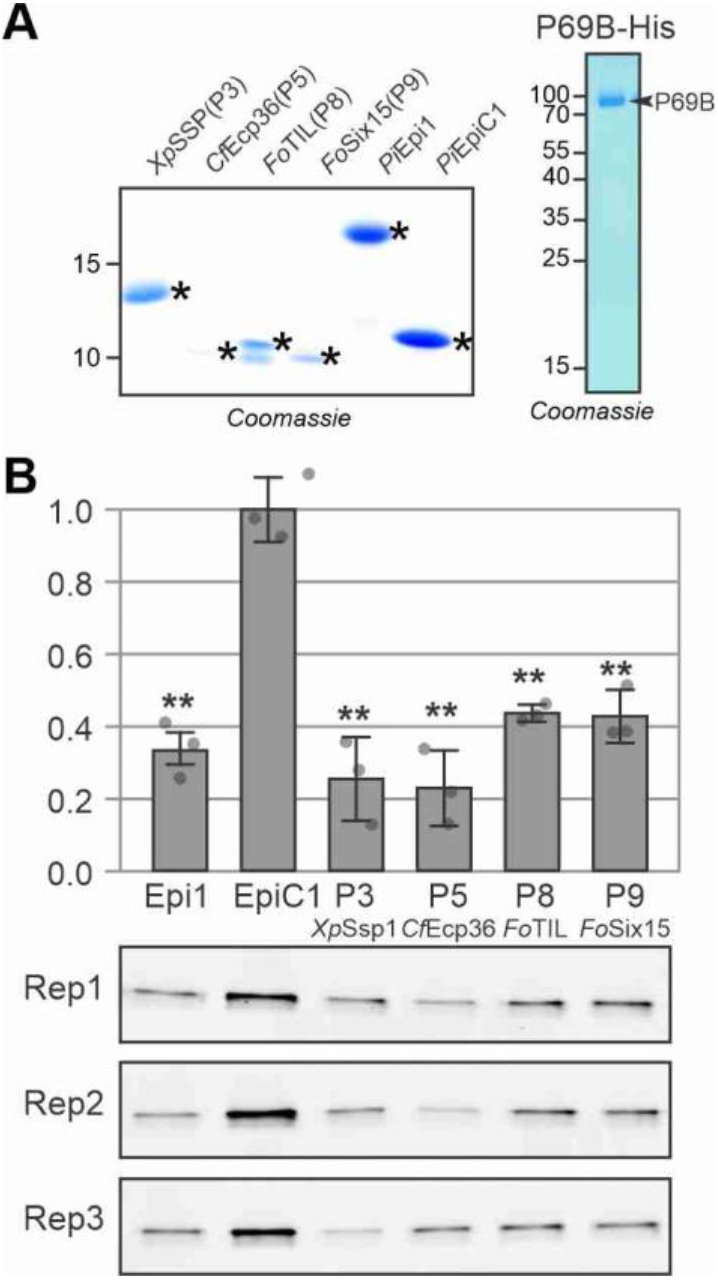
Activity labelling of P69B is suppressed by four novel inhibitors **(A)** Purified candidate inhibitors. Candidate inhibitors and Epi1a (positive control) and EpiC1 (negative control) were expressed in *E. coli* as fusion proteins with N-terminal His-MBP-TEV. The fusion proteins were purified over Ni-NTA and amylose resin, subsequently, and then cleaved by TEV protease. See Supplemental **Figure S1** for the full gel. His-TEV protease and purification tags were removed using Ni-NTA and MW cut off filter and SSPs were used for inhibition assays in (C). **(B)** P69B-His was transiently expressed in *Nicotiana benthamiana* by agroinfiltration (Paulus et al., 2020) and purified over Ni-NTA from apoplastic fluids isolated at 5 days-post-agroinfiltration. The eluate was analysed on protein gel stained with Coomassie (shown here) and used for inhibition assays (in (C)). **(C)** All four candidate inhibitors and the Epi1a but not EpiC1 suppress activity-based labelling of P69B with FP-TAMRA. Purified P69B-His was pre-incubated with purified (candidate) inhibitors at a 1:100 molar ratio and then labelled with FP-TAMRA in n=3 replicates. Proteins were separated on protein gel and scanned for fluorescence. Fluorescence was quantified and the signal intensity of the negative control (EpiC1) was set at 100% labelling to calculate the relative labelling upon preincubation with the positive control (Epi1a) and the four candidate inhibitors. Error bars represent STDEV of n=3 replicates. **, p<0.01 (p-values from two-sided, pairwise t-tests were adjusted for multiple testing using the Benjamini-Hochberg procedure)

To test for P69B inhibition, the purified inhibitor candidates and the Epi1 and EpiC1 controls were preincubated with purified P69B. Subsequent labelling with FP-TAMRA and detection from protein gels by fluorescence scanning revealed that P69B labelling is significantly reduced upon preincubation with Epi1 and all four novel candidate inhibitors, when compared to the EpiC1 negative control (**Figure 4B**). These data confirm that all four tested novel candidate inhibitors indeed inhibit P69B.

### P69B is an effector hub targeted by five distinct pathogen-derived inhibitors

We finally investigated the four novel P69B inhibitors more closely, by studying their AFM-predicted binding to P69B in combination with alignments of inhibitor homologs from public databases (**Figure 5**). Mapping sequencing reads from eleven wild tomato species against the tomato reference genome to generate phased P69B alleles from wild tomato relatives revealed that P69B has only one hyper-variant residue at position 400, being either His, Arg, Asp or Gly (Supplemental **Figure S3**). Interestingly, this variant site locates close to the substrate binding groove in P69B (**Figure 5A**).

**Figure 5.**
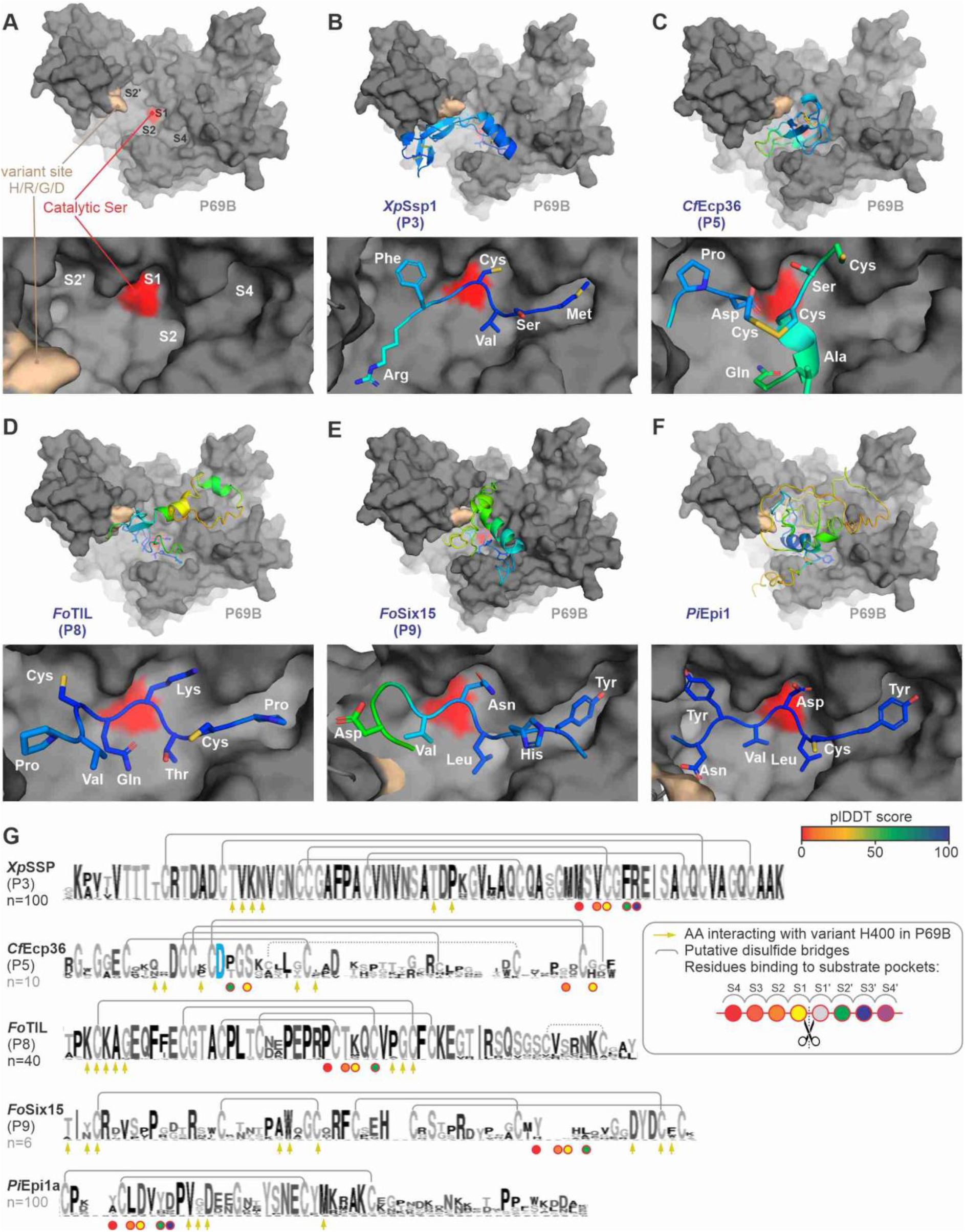
P69B is an effector hub targeted by five pathogen-derived inhibitors. AFM-predicted models of P69B without inhibitor **(A)**, or with *Xp*Ssp1(P3); **(B)** *Cf*Ecp36(P5); **(C)** *Fo*TIL(P8); **(D)** *Fo*Six15(P9) **(E)** and *Pi*Epi1 **(F)**. P69B is shown in a gray surface representation with variant residues (crème) and the active site (red), in the substrate binding groove that has substrate binding pockets (S4-S2-S1-S2’) that bind to substrate/inhibitor residues P4, P2, P1 and P2’, respectively. The inhibitors are shown as cartoon and sticks and coloured using a rainbow scheme based on their plDDT scores, which range from 0 (worst) to 100 (best). The zoomed image (bottom) shows the predicted occupation of the substrate binding pockets in P69B by different residues of the inhibitor. **(G)** Sequence conservation between homologs of the identified P69B inhibitors. Shown is the sequence logo for n=x close homologs from other plant pathogen species, identified by BLAST searches in the NCBI database and presented in Weblogo (Crooks et al., 2004). Highlighted are the residues that probably interact with the substrate binding pockets in P69B (circles); the conserved Asp residue in *Cf*Ecp36 that interacts with the catalytic site (blue); the residues that might interact with the variant residue in P69B (arrows); and putative disulphide brides observed in the AFM model (grey lines).

The first newly discovered P69B inhibitor is an SSP of the bacterial tomato pathogen *Xanthomonas perforans* named *Xp*Ssp1. *Xp*Ssp1 fits nicely in the substrate binding groove of P69B with high plDDT scores at the interface (**Figure 5B**). *Xp*Ssp1 is highly conserved in plant pathogenic *Xanthomonas* species and contains five conserved disulphide bridges and several residues that contact the hypervariable residue in P69B (**Figures 5G**). A conserved methionine, valine, and phenylalanine occupy the S4, S2 and S2’ pockets in P69B (**Figure 5B**). And a conserved disulphide bridge occupies the S1 pocket and this structure is probably the reason why this SSP inhibits P69B. The *Xp*Ssp1 ortholog in *Xanthomonas oryzae* pv. *oryzicola* (XOC_0943) is expressed during infection of rice (Liao et al., 2019), so it is likely that *Xp*Ssp1 homologs play an active role during *Xanthomonas* infections.

The second newly discovered P69B inhibitor is from the fungal pathogen *Cladosporium fulvum* and has been previously detected in apoplastic fluids from infected plants by proteomics as Extracellular Protein-36 (*Cf*Ecp36, Mesarich et al., 2018). Its detection by proteomics is consistent with a high expression of the *CfEcp36* gene during infection: 479 tpm in infected tissues. The binding of *Cf*Ecp36 is distinct from all the other inhibitors as it does not use a single strand to occupy the substrate binding groove (**Figure 5C**). Instead, *Cf*Ecp36 uses two strands and probably utilizes two disulphide bonds and an aspartate interacting with two active site residues to avoid processing by P69B (**Figure 5C**). *Cf*Ecp36 has homologs in other ascomycete plant pathogens including *Zymoseptoria, Verticillium* and *Colletotrichum* that share the aspartate and five AFM-predicted disulphide bridges (**Figure 5G**). Several variant residues in *Cf*Ecp36 homologs are in close proximity to the hyper-variant residue in P69B (**Figures 5C** and **5G**).

Two newly discovered P69B inhibitors are from the fungal pathogen *Fusarium oxysporum*. Both are highly expressed during infection, reaching 683 and 717 tpm in infected tissues. The first P69B inhibitor has homology to a trypsin-inhibitor-like protein (Rosengren et al., 2001), and is hence coined *Fo*TIL. Although the overall structure of *Fo*TIL has an intermediate confidence, *Fo*TIL binds with high plDDT scores to the substrate binding grove occupying S4, S2, S1 and S2’ pockets with proline, threonine, lysine and cysteine residues, respectively (**Figure 5D**). The cysteine residues at the P3 and P2’ positions are involved in predicted disulphide bridges that probably constrain the structure so it remains uncleaved by P69B. *Fo*TIL has close homologs in many *Fusarium* species and shares high homology that includes four of the five putative disulphide bridges and conserved residues that might interact with the hyper-variant residue in P69B (**Figure 5G**). Interestingly, although these proteins are highly conserved, the residue predicted to occupy the S1 pocket is highly variant (K, Q, M or D).

The other P69B inhibitor of *Fo* has previously been described as secreted-into-xylem protein-15 (*Fo*Six15, Simbaqueba et al., 2021). *Fo*Six15 uses a strand to occupy the S4, S2 and S1 pockets in P69B with tyrosine, leucine and asparagine residues with high confidence (**Figure 5E**). *Fo*Six15 has homologs in fungal plant pathogens *Dactylonectria* and *Ramularia* that share four highly conserved disulphide bridges and are otherwise highly polymorphic, including the residues that occupy the S4-S2-S1 pockets, though some of the residues that might interact with the hyper-variable residue in P69B seem more conserved (**Figure 5G**).

These four novel P69B inhibitors are structurally distinct from each other and from the previously described Kazal-like *Pi*Epi1, which is predicted to occupy the S4, S2 and S1 and S2’ pockets using tyrosine, leucine, aspartate and tyrosine residues, respectively (**Figure 5F**). Epi1 has many homologs in plant pathogenic *Phytophthora* species that share two disulphide bridges, but residues are more polymorphic at positions that are predicted to occupying S1 and S2 pockets or interact with the hypervariable residue in P69B (**Figure 5G**). Overall, despite the structural diversity, most P69B inhibitors seem to occupy the S4 and S2 pockets with similar residues but the predicted residues occupying the S1 pocket can be strikingly diverse and include both basic (Lys) and acidic (Asp) residues, as well as serine, asparagine and a disulphide bridge.

## DISCUSSION

We successfully used AFM as a discovery tool to identify novel cross-kingdom interactions at the plant-pathogen interface. We used AFM to predict complexes between 1,879 SSPs with six extracellular hydrolases and from 376 complexes with high scores, we manually selected 15 putative inhibitors that block the active site with an intrinsic fold and are likely expressed during infection. Four of the candidates were expressed and confirmed to be P69B inhibitors. This work demonstrates that the use of artificial intelligence to predict cross-kingdom protein complexes can make instrumental contributions to host-microbe interactions.

We found that the vast majority of the SSPs in AFM-predicted complexes with high scores are probably not hydrolase inhibitors. Some might, however, rather be substrates or allosteric inhibitors, which remains to be explored in the future. We could, however, successfully identify novel inhibitor candidates because we used a stringent selection by manually screening the structures for SSPs that block the active site and have an intrinsic structure. This selection resulted in a high hit rate because all four tested candidates were confirmed to be P69B inhibitors.

In addition to previously described Kazal-like inhibitors of *Phytophthora infestans*, we discovered four novel P69B inhibitors from three additional tomato pathogens: *Xp*Ssp1 from *Xanthomonas perforans; Cf*Ecp36 from *Cladosporium fulvum* and *Fo*TIL and *Fo*Six15 from *Fusarium oxysporum*. These pathogens secrete P69B inhibitors because they are exposed to very high levels of P69B during apoplast colonisation. This suggests that other tomato pathogens probably also secrete P69B inhibitors that remain to be identified. We may have missed some putative P69B inhibitors produced by other pathogens because they were too large (>35 kDa), were not predicted to be secreted, were not detected in the used transcriptomic dataset, were false negatives in AFM modeling, or are not proteinaceous in nature.

Our AFM screen also uncovered seven inhibitor candidates of chitinases, which remain to be validated experimentally. Pathogen-secreted inhibitors of chitinases were not reported before but are likely to exist. The existence of Class-I chitinase inhibitors was implicated by the accumulation of variant residues around the substrate binding groove (Bishop et al., 2000). Interestingly, in our AFM-predicted complexes, these variant positions might directly interact with the predicted inhibitors of *Ralstonia solanacearum* and *Fusarium oxysporum* (Supplemental **Figure S4**). It might be counterintuitive that also bacteria secrete putative chitinase inhibitors even though they do not have chitin in their cell wall. However, chitinases may have alternate activities. LYS1, for instance, belongs to the Class-III chitinases but hydrolyses peptidoglycan in the bacterial cell wall (Liu et al., 2014), and *Nb*PR3 belongs to the Class-II chitinases but has antibacterial activity and no chitinase activity (Sueldo et al., 2023). It seems likely that other proclaimed chitinases may have antibacterial activities and that this is why they are targeted by bacterial inhibitors.

The fact that P69B is targeted by so many pathogens indicates that it plays an important role in immunity against different pathogens. So far, immunity phenotypes upon P69B depletion remain to be described. P69B is, however, required for the activation of immune protease Rcr3 (Paulus et al., 2020) and for processing the *Pi*-secreted SSP PC2, which then triggers the hypersensitive response (HR, Wang et al., 2021). It seems likely that P69B has many additional substrates in tomato and its apoplastic pathogens. Interestingly, our AFM screen identified 17 pathogen-produced SSPs that interact with the substrate binding groove of P69B and might be substrates that can be studied further.

P69B inhibition is associated with diversification in two directions. At the species level, we detected polymorphism within P69B orthologs at position 400. The AFM models suggest that this residue might directly interfere with P69B inhibitors. In addition to the selection pressure on P69B orthologs, the selection probably also resulted in the diversification of *P69* paralogs in *Solanum* species. There are nine *P69B* paralogs in tomato and all these 10 genes (*P69A-J*) form a gene array at a single genomic cluster on chromosome 8 (Supplemental **Figure S5A**). These *P69B* paralogs are all inducible by biotic stress but their transcriptional induction varies between cultivars and pathogen (Supplemental **Figure S5B**). Interestingly, residue variation between P69 paralogs mostly locates to the edge of the substrate binding groove (Supplemental **Figure S5C**). These ‘ring-of-fire’ positions will likely cause differential sensitivity of the paralogs for the different pathogen-derived inhibitors. This variation indicates that the P69B paralogs evolved from parallel arms races with pathogen-secreted inhibitors, resulting in gene duplication and diversification in the ancestral *Solanum* species. Taken together, these observations indicate a fascinating new arms race at the plant-pathogen interface.

Although we report a successful use of AFM in predicting novel cross-kingdom interactions, we did notice that AFM can produce false negative scores. Some well-established inhibitor-hydrolase interactions receive relatively low ipTM+pTM scores. Avr2-Rcr3 for instance, scored only 0.44, despite being well-established (Rooney et al., 2005). Scores were also unexpectedly low for Vap1-Rcr3 (0.51); SDE1-RD21a (0.53), Pit2-CP1A (0.35), Pep1-Pox12 (0.37), and Gip1-EGase (0.28), despite their reported interactions (Lozano-Torrez et al., 2012; Clark et al., 2018; Mueller et al., 2013; Hemetsberger et al., 2013; Damasenco et al., 2008). These low scores indicate that our AFM screen can result in false negatives. Some of the low scores might be due to low mean non-gap MSA depth for some of the SSPs, which is below the desired 100 MSA for 45% of the tested SSPs. This implies that new interactions might be discovered when additional SSP sequences are added to the database.

The simultaneous discovery of four novel P69B inhibitors demonstrate that artificial intelligence can be a powerful ally in the prediction of novel cross-kingdom interactions at the plant-pathogen interface. This *in-silico* interactomic approach overcomes important limitations of traditional assays such as Y2H, CoIP and phage display, which are challenging to apply for secreted proteins having disulphide bridges and interacting at apoplastic pH (pH 5-6). Some of the current limitations of AFM might be overcome by increased sequencing and by quickly by improving prediction algorithms, evaluation and verification methods such as AF2Complex (Gao et al., 2022), AlphaPulldown (Ye et al., 2023) and PAE viewer (Elfmann & Stülke, 2023). Screens for hydrolase inhibitors can be automated using a script that searches for residues of candidate inhibitors that are in close proximity to the active site. Certainly, artificial intelligence to predict plant-pathogen interactions will be a revolutionary new approach in future research.

## MATERIALS AND METHODS

### Protein complex prediction with AFM

Protein complexes were modelled using AlphaFold multimer v2.1.1 (Jumper et al. 2021; Evans et al. 2021). Template sequence searches of individual proteins were re-used to model protein complexes as they are identical between Alphafold2 and AFM. The AFM-specific database search against the unclustered Uniprot database with JackHMMer v3.3 was added for each monomer as in AFM (Supplemental **File S1**, script-1) For each protein complex, AFM additionally matched hidden Markov models extracted from the Uniref90 MSA against the PDB seqres database. The small bfd database was used and all databases were downloaded as instructed in the ‘download_all_data.sh’ file from the AlphaFold2 v2.1.1 release on GitHub. The sequences for the four control complexes are in Supplemental **File S2**. The structure files (.pdb) of the four control complexes and 15 putative inhibitor-hydrolase complexes are provided in Supplemental **File S3**.

### Analysing output parameters of AFM

Mean non-gap amino acid depth for chains of each protein were calculated using the features.pkl output file generated by AFM (Supplemental **File S1**, script-2). Mean non-gap MSA depths for proteins modelled in several different complexes are the mean of their mean non-gap MSA depths from all complexes. Total computing time calculations of AFM were based on the timings.json file of each protein complex. To calculate CPU and GPU hours based the timings.json files, it is necessary to know that all AlphaFold2 monomer computations were completed with eight CPU cores and one GPU at any time. AFM computations were executed with one CPU core and one GPU at any time.

### Tomato and plant pathogen proteomes and transcriptomes

Amino acid sequences of tomato proteins were from the *S. lycopersicum* ITAG4.0 proteome (Tomato Genome Consortium, 2012). Tomato amino acid sequences of Solyc09g098540.3.1 (class I chitinase), Solyc05g050130.4.1 (class III chitinase), Solyc07g005090.4.1 (class V chitinase), Solyc08g079870.3.1 (P69B), Solyc02g077040.4.1 (Pip1) and Solyc08g067100.2.1 (AED1) are listed in Supplemental **File S4**. The *P. syringae* pv. tomato DC3000 proteome (https://pseudomonas.com/downloads/pseudomonas/pgd_r_20_2/Pseudomonas_syringae_pv_tomato_DC3000_111/Pseudomonas_syringae_pv_tomato_DC3000_111.faa.gz) and transcriptome (https://pseudomonas.com/downloads/pseudomonas/pgd_r_20_2/Pseudomonas_syringae_pv_tomato_DC3000_111/Pseudomonas_syringae_pv_tomato_DC3000_111.ffn.gz) were from the GCF_000007805.1 genome assembly, the *X. perforans* DMS 18975 proteome (https://ftp.ncbi.nlm.nih.gov/genomes/all/GCF/013/112/235/GCF_013112235.1_ASM1311223v1/GCF_013112235.1_ASM1311223v1_protein.faa.gz) and transcriptome (https://ftp.ncbi.nlm.nih.gov/genomes/all/GCF/013/112/235/GCF_013112235.1_ASM1311223v1/GCF_013112235.1_ASM1311223v1_cds_from_genomic.fna.gz) were from the GCF_000009125.1 genome assembly, the *R. solanacearum* GMI1000 proteome (https://ftp.ncbi.nlm.nih.gov/genomes/all/GCF/000/009/125/GCF_000009125.1_ASM912v1/GCF_000009125.1_ASM912v1_protein.faa.gz) and transcriptome (https://ftp.ncbi.nlm.nih.gov/genomes/all/GCF/000/009/125/GCF_000009125.1_ASM912v1/GCF_000009125.1_ASM912v1_cds_from_genomic.fna.gz) were from the GCF_000009125.1 genome assembly, the *B. cinerea* B05.10 proteome (https://ftp.ncbi.nlm.nih.gov/genomes/all/GCF/000/143/535/GCF_000143535.2_ASM14353v4/GCF_000143535.2_ASM14353v4_protein.faa.gz) and transcriptome (https://ftp.ncbi.nlm.nih.gov/genomes/all/GCF/000/143/535/GCF_000143535.2_ASM14353v4/GCF_000143535.2_ASM14353v4_rna.fna.gz) were from the GCF_000143535.2 genome assembly, the *F. oxysporum* f. sp. *lycopersici* 4287 proteome (https://ftp.ncbi.nlm.nih.gov/genomes/all/GCF/000/149/955/GCF_000149955.1_ASM14995v2/GCF_000149955.1_ASM14995v2_protein.faa.gz) and transcriptome (https://ftp.ncbi.nlm.nih.gov/genomes/all/GCF/000/149/955/GCF_000149955.1_ASM14995v2/GCF_000149955.1_ASM14995v2_rna.fna.gz) were from the GCF_000149955.1 genome assembly, the *C. vulvum* Race5_Kim proteome (https://ftp.ncbi.nlm.nih.gov/genomes/all/GCA/020/509/005/GCA_020509005.1_Cfulv_R5_v4/GCA_020509005.1_Cfulv_R5_v4_protein.faa.gz) and transcriptome (https://ftp.ncbi.nlm.nih.gov/genomes/all/GCA/020/509/005/GCA_020509005.1_Cfulv_R5_v4/GCA_020509005.1_Cfulv_R5_v4_cds_from_genomic.fna.gz) were from the GCA_020509005.1 genome assembly, the *P. infestans* T30-4 proteome (https://ftp.ncbi.nlm.nih.gov/genomes/all/GCF/000/142/945/GCF_000142945.1_ASM14294v1/GCF_000142945.1_ASM14294v1_protein.faa.gz) was from the GCF_000142945.1 genome assembly.

### Prediction of small secreted proteins (SSPs)

A custom secretion prediction pipeline was used to predict small secreted proteins likely to remain in the apoplast (Homma, 2022). Proteins were considered apoplastic proteins if they were predicted to be secreted by either SignalP5.0 or TargetP2.0 or both and were predicted to be localised in the apoplast by ApoplastP1.0.1. Proteins were considered small if their full-length sequence was predicted to <35 kDa. If a protein had been predicted by SignalP5.0 to be secreted, we used the mature sequence as predicted by SignalP5.0. If a sequence was only predicted by TargetP2.0 to be secreted, the mature sequence as predicted by TargetP2.0. An additional 14 known apoplastic proteins were added from *C. fulvum* and *F. oxysporum* f. sp. *lycopersici* that did not have identical copies in the predicted proteomes used for this study. These additional 14 proteins included *C. fulvum* proteins AIZ11404.1 (Avr2), AHY02126.1 (Avr5) and AQA29222.1 (Ecp17) and *F.oxypsorum* f. sp. *lycopersici* proteins ALI88770.1 (Six1), UEC48541.1 (partial Six3), BAM37635.1 (Six4), ALI88836.1 (Six6), AIY35187.1 (Six7), ACN69118.1 (Six8), AGG54051.1 (Six10), AGG54052.1 (Six11), ANF89367.1 (Six12), AGG54055.1 (Six14) and APP91304.1 (Six15). All mature, small, putatively apoplastic pathogen-derived proteins were filtered against any duplicated amino acid sequences using seqkit (Shen et al. 2016). All mature 1,879 SSP sequences used for the AFM screen are in Supplemental **File S5**.

### RNA-seq data mining, raw reads filtering and mapping of trimmed reads

Publicly available raw-read RNA-seq data sets were downloaded of infected plant tissue for *R. solanacearum* infecting tomato petioles (SRR5467166, SRR5467167, SRR5467168), *B. cinerea* infecting tomato leaves (SRR6924534, SRR6924535, SRR6924536), *F. oxysporum* f. sp. l*ycopersici* infecting tomato roots (SRR6050413, SRR6050414) and *C. fulvum* infecting tomato leaves (SRR1171035, SRR1171040, SRR1171043, SRR1171047) from NCBI’s sequence read archive. No suitable *in planta* RNA-seq dataset for *X. perforans* was identified. Each sequencing read was labeled by its likely source of origin with Centrifuge 1.0.4 (Kim et al., 2016) using the NCBI nucleotide non-redundant sequences, last updated 03/03/2018. Putative host-derived RNA reads were removed by filtering against taxonomic ids 3700 (Brassicaceae), 3701 (Arabidopsis), 3702 (*A. thaliana*), 4070 (Solanaceae), 4081 (*S. lycopersicum*) and 4107 (Solanum). Filtered RNA-seq reads were quality trimmed using timmomatic 0.39 (‘LEADING:3 TRAILING:3 SLIDINGWINDOW:4:15 MINLEN:36’ for unpaired and paired-end reads) (Bolger et al. 2014). Host-filtered and quality-trimmed reads were mapped onto nucleotide sequences of the respective plant pathogen using Kallisto v0.46.2 (Bray et al. 2016). Genes were considered expressed during infection if they exceed an average gene expression ≥ 2 transcripts per million (tpm). The minimum expression level of EBI’s gene expression atlas is 0.5 tpm.

### Generating sequences of P69B orthologs in wild tomato species

Publicly available genomic sequencing reads of eleven wild tomato species from NCBI’s sequence read archive were downloaded: *S. lycopersicum* var. c*erasiforme* BGV006865 (SRR7279628), *S*. *pimpinellifolium* LA2093 (SRR12039813), *S*. *cheesmaniae* LA0483 (ERR418087), *S. arcanum* LA2157 (ERR418092), *S. neorickii* LA2133 (ERR418090), *S. hualylasense* LA1983 (ERR418095), *S. chilense* LA3111 (SRR13259416), *S. corneliomuelleri* LA0118 (ERR418061), *S. peruvianum* LA1954 (ERR418094), *S. habrochaites* LYC4 (ERR410237) and *S. pennellii* LA0716 (ERR418107) (100 Tomato Genome Sequencing Consortium 2014; Stam et al., 2019; Wang et al., 2020). Genomic sequencing reads were quality trimmed using trimmomatic v0.39 with the following settings ‘LEADING:3 TRAILING:3 SLIDINGWIN-DOW4:15 MINLEN:36’ (Bolger et al. 2014). Reads were mapped against the Sol4.0 *S. lycopersicum* reference genome assembly using BWA-MEM v0.7.17 (Li 2013). Mapped reads were processed and sorted using Samtools v1.7 (Li et al. 2009; Li 2011). InDels were realigned using GATK v3.8-1-0-gf15c1c3ef (McKenna et al., 2010). Variants were called using bcftools v1.7 using a phred score of 20 as a cut off (Li 2011). And phased using whatshap v1.0 (Martin et al. 2016). Coding sequences from different species were generated from loci using exonerate v2.4.0 (Slater and Birney 2005). These alleles were generated using three standardised snakemake v6.7.0 workflows (Homma 2022b, c, d).

### P69B cloning and purification

First, pJK187 was generated by introducing fragments from pAGM4723, pICH41308, pICH51288 and pICH41414 (Weber et al., 2011; Engler et al., 2014) into pJK001 (Paulus et al., 2020), resulting in a binary plasmid that contains the 35S promoter and 35S terminator with the *nptII* kanamycin and LacZ as the fragment to be replaced by insert sequences.

The gene sequence of P69B (with *Nt*PR1a signal peptide, see Supplemental **Table S1**) was synthesized at Twist Bioscience and inserted into the binary vector pJK187 using BpiI to yield *Nt*PR1a-P69B-His (pFH20). Plasmids were sequenced using Source Bioscience using 35S promoter (CTATCCTTCGCAAGACCCTTC) and terminator (CTCAACACATGAGCGAAACC) primers to confirm the inserts. Validated binary plasmids were transformed into *A. tumefaciens* GV3101 (pMP90) via heat shock transformation.

Four-week-old *N. benthamiana* plants were infiltrated with a 1:1 mixture of *Agrobacterium tumefaciens* GV3101(pMP90) OD_600_ = 0.5) containing pFH20 and silencing suppressor p19 (Van der Hoorn et al., 2003), respectively. Apoplastic fluid containing P69B-His was extracted 5 days after infiltration as previously described (Kourelis et al., 2020). The recombinant protein of P69B-His was purified by HisPur™ Ni-NTA resin and concentrated in 25 mM Tris-HCl pH=6.8 using a 50 kDa MWCO Amicon Ultra-15 filter.

### Expression and purification of putative inhibitors

A sequence encoding His-MBP-TEV was synthesized at Twist Bioscience (South San Francisco, Supplemental **Table S1**) and inserted into the pET-32/28 vector (Novinec et al., 2012) using NheI and XhoI restriction sites to generate the pET-32/28-His-MBP-TEV vector pHJ000 (Supplemental **Table S2**). Codon-optimized sequences encoding the different candidate inhibitors were synthesized at Twist Bioscience (Supplemental **Table S1**), amplified using cloning primers (Supplemental **Table S3**) and ligated into the pHJ000 using ClonExpress Ultra One Step Cloning Kit (Vazyme Biotech) to yield His-MBP-inhibitor constructs pHJ028 (P3, *Xp*Ssp1); pHJ033 (P5, *Cf*Ecp36); pHJ029 (P6); pHJ032 (P7); pHJ030 (P8, *Fo*TIL); pHJ031 (P9, *Fo*Six15), respectively (Supplemental **Table S2**). The gene fragments of Epi1 and EpiC1 were amplified from pFlag-Epi1 (Tian et al., 2004) and pJK155 (pET28b-T7::OmpA-HIS-TEV-EpiC1), respectively, to yield constructs pHJ046 (*Pi*Epi1) and pHJ047 (*Pi*EpiC1), respectively. All the cloning primers and sequencing primers are provided in Supplemental **Table S3**.

The plasmids were transformed into *E. coli* Rosetta-gami B(DE3)pLysS (Novagen, Sigma-Aldrich) and cultures in LB (Luria-Bertani) liquid medium were induced with 0.1 mM isopropyl β-D-1-thiogalactopyranoside (IPTG) and incubated at 18°C for 24 h. Cells were pelleted by centrifugation at 8,000 x *g* for 5 min and the supernatant was discarded. The cell pellet was resuspended in 50 mM Tris-HCl, pH 7.5. The CelLytic™ Express (Sigma-Aldrich) was used for bacterial cell lysis, and the supernatant was collected for further protein purification. The recombinant proteins were purified using HisPur™ Ni-NTA resin (Thermo Fisher Scientific) and amylose resin (NEB), and then the tobacco etch virus (TEV) protease (Sigma-Aldrich) was added to remove the purification tags. His-tagged TEV protease and purification tags were removed over Ni-NTA and a 30 kDa Amicon filter, whilst concentrating the cleaved inhibitor protein in 25 mM Tris-HCl pH 6.8. Inhibitors were used immediately or stored at −80°C.

### Inhibition assays

The Bio-Rad DC Protein assay kit was used to measure the protein concentration of candidate inhibitors and P69B. To test the P69B inhibition, 85 pmol purified candidate inhibitors were preincubated with 0.85 pmol purified P69B-His protein at room temperature for 0.5 hours in 25 mM Tris-HCl (pH 6.8), 1 mM DTT, and then labelled by adding 0.5 μM FP-TAMRA (Thermo-Fisher) and incubating for 1 hr at room temperature in the dark. The labelling reaction was stopped by adding 4× loading buffer (200 mM Tris-HCl (pH 6.8), 400 mM DTT, 8% SDS, 0.2%bromophenol blue, 40% glycerol) and boiling for 7 min. at 95 °C. Samples were separated on 15% SDS-PAGE gel. The gel was washed three times with Milli-Q water and scanned for fluorescence with the Typhoon scanner (GE Healthcare) using a Cy3 setting. Signal intensities were quantified using ImageJ and normalized to the EpiC1 negative control. Statistical testing of inhibition was based on two-sided, pairwise comparisons between the putative inhibitor and the EpiC1 negative control. Calculated p-values were adjusted for multiple testing using the Benjamini-Hochber procedure.

## Supporting information

Suppplemental File S2

## DECLARATIONS

### Availability of data and materials

All data generated or analysed during this study are included in this article and its supplementary information files.

### Authors’ contributions

FH and RH conceived the project; FH performed all bioinformatic analysis; JH produced candidate inhibitors and P69B and performed inhibition experiments; RH wrote the manuscript with input from all authors. The funding body had no influence on the design of the study and collection, analysis, and interpretation of data and in writing the manuscript.

## Acknowledgements

We like to thank Urszula Pyzio for excellent plant care, Sarah Rodgers and Caroline O’Brien for excellent technical support; Dr. Jiorgos Kourelis for constructing pJK187; Dr. Sheng Huang (Guangxi University, Nanning Guangxi, China) for providing expression data of XOC_0943; Dr. Brian Mooney, Dr. Mariana Schuster, and Dr. Nattapong Sanguankiattichai for excellent suggestions and the Advanced Research Computing (ARC) facility of the University of Oxford for continuous access to their high-performance computing cluster.

## Supplemental Materials

**Supplemental Figure S1** Production of candidate inhibitors and P69B for inhibition assays.

**Supplemental Figure S2** MSA depth of SSPs and hydrolases used in Figure 2.

**Supplemental Figure S3** Amino acid alignment of P69B orthologs from wild tomato species. **Supplemental Figure S4** Two candidate inhibitors interact with positively selected sites in Class-I chitinases.

**Supplemental Figure S5** Expression and variation of the ten P69 paralogs of tomato.

**Supplemental File S1** Used scripts (.zip file)

**Supplemental File S2** Protein sequences of inhibitors and proteases used for Figure 1 (.doc)

**Supplemental File S3** PDB files of 5 controls and 15 candidate complexes (.zip)

**Supplemental File S4** Sequences of six hydrolases used for AFM screen (.doc)

**Supplemental File S5** sequences of mature SSPs used for AFM screen (.fasta)

**Supplemental Table S1** Synthesised nucleotide sequences (.doc)

**Supplemental Table S2** Used plasmids (.doc)

**Supplemental Table S3** Primer sequences (.doc)

