## Supplemental Figure for "Alphafold-multimer predicts cross-kingdom interactions at the plant-pathogen interface"

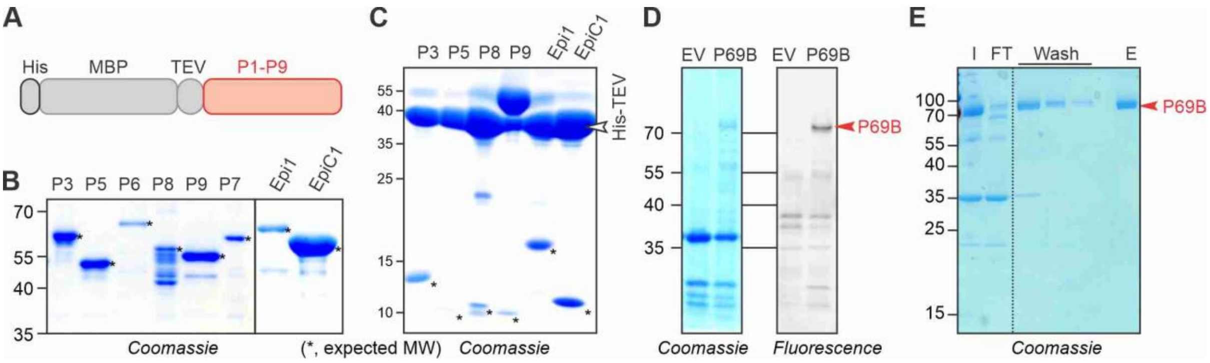


**Figure S1** Production of candidate inhibitors and P69B for inhibition assays.

**(A)** Candidate inhibitors lacking their endogenous signal peptides were codon-optimized for expression in *E. coli* and cloned in-frame with a 6-His purification tag; the maltose binding protein (MBP) and the cleavage site of tobacco etch virus (TEV) protease. **(B)** Fusion proteins were expressed in *E. coli* and purified on HisPur™ Ni-NTA resin and amylose resin, subsequently. Fusion proteins were detected at their predicted molecular weight (MW, *) by SDS-PAGE and Coomassie staining. **(C)** Putative inhibitors were released from their purification tags upon incubation with His-TEV protease. **(D)** P69B-His is efficiently expressed by agroinfiltration. P69B with a C-terminal His tag and an empty vector (EV) control were transiently expressed by agroinfiltration. Apoplastic fluids were isolated, labeled with FP-TAMRA, separated on SDS-PAGE and scanned for fluorescence (right) and stained with Coomassie (left). **(E)** Purification of P69B-His on Ni-NTA. I, input sample; FT, flow through; E, eluate.

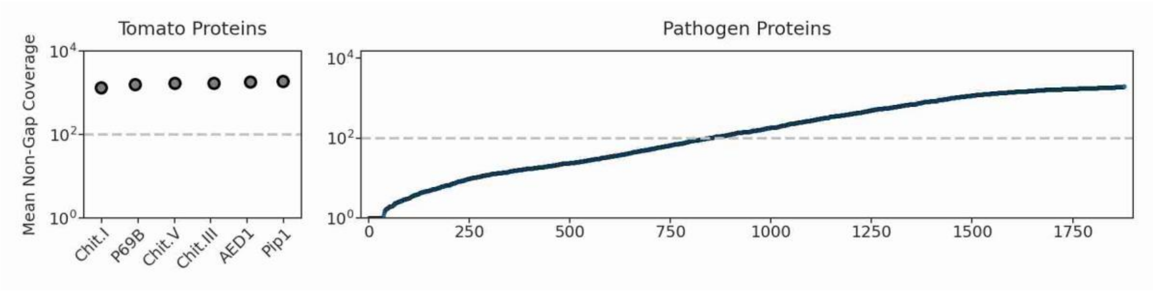


**Figure S2** MSA depth of SSPs and hydrolases used in Figure 2.

The mean non-gap multiple sequence alignment (MSA) depth is shown for the tomato hydrolases (left) and the 1,879 SSPs (right), ranked by MSA depth. The dash line indicates the desired minimum of 100 MSA depth.

```
      1                                                                                                  100
lycop MGLLKILLVF IFCSFQWPTI QSNLETYIVH VESPESLVTT QSLLTDLGSY YLSFLPKTAT TISSSGNEEA ATMIYSYHNV MTGFAARLTA EQVKEMEKKH
pimpi MGLLKILLVF IFCSFQWPTI QSNLETYIVH VESPESLVTT QSSLTDLGSY YLSFLPKTAT TISSSGNEEA ATMIYSYHNV MTGFAARLTA EQVKEMEKKH
chees MGLLKILLVF IFCSFQWPTI QSNLETYIVH VESPESLVTT QSSLTDLGSY YLSFLPKTAT TISSSGNEEA ATMIYSYHNV MTGFAARLTA EQVKEMEKKH
habro MGLLKILLVF IFCSFPCPTI QSNFETYIVH VESPESLVTT LSSLTDLDSY YLSFLPKTAT TISSSGNEEA ATMIYSYHNV MTGFAARLTA EQVKEMEKKH
corne MGLLKILLVF IFCSFPWSTI QSNLETYIVH VESPESLVTT QSSLTDLDSY YLSFLPKTAT IISSSGNEEA ATMIYSYHNV MTGFAARLTA EQVKEMEKKH
neori MGLLKILLVF IFCSFPWPTI QSNLETYIVH VESPESLVTT QSSLTDLDSY YLSFLPKTAT TISSSGNEEA ATMIYSYHNV MTGFAARLTA EQVKEMEKKH
arcan MGLLKILLVF IFCSFQWPTI QSNLETYIVH VESPESLVTT QSSLTDLDSY YLSFLPKTAT TISSSGNEEA ATMIYSYHNV MTGFAARLTA EQVKEMEKKH
huayl MGLLKILLVF IFCSFQWPSI QSNLETYIVH VESPESLVTT QSSLTDLDSY YLSFLPKTAT TISSSGNEEA ATMIYSYHNV MTGFAARLTA EQVKEMEKKH
chile MGLLKILLVF IFCSFPWTTI QSNLETYIIH VESPESLVTT QSSLTDLDSY YLSFLPKTTT TISSSGNEEA ATMIYSYHNV MTGFAARLTA EQVKEMEKKH
penne MGLLKILLVF IFCSFPWPTI QSNLETYIVH VESPESLVTT QSSLTDLDSY YLSFLPKTTT TISSSGNEEA ATMIYSYHNV MTGFAARLTA EQVKEMEKKH
peruv MGLLKILLVF IFCSFPWPTI QSNLETYIVH VESPESLVTI QSSLTDLDSY YLSFLPKTTT TISSRGNEEA ATMIYSYHNV MTGFAARLTA EQVKEMEKKH

      101                                                                                                200
lycop GFVSAQKQRI LSLHTTHTPS FLGLQQNMGV WKDSNYGKGV IIGVIDTGII PDHPSFSDVG MPPPPAKWKG VCESNFTNKC NNKLIGARSY QLGNGSPIDS
pimpi GFVSAQKQRI LSLHTTHTPS FLGLQQNMGV WKDSNYGKGV IIGVIDTGII PDHPSFSDVG MPPPPAKWKG VCESNFTNKC NNKLIGARSY QLGNGSPIDS
chees GFVSAQKQRI LSLHTTHTPS FLGLQQNMGI WKDSNYGKGV IIGVIDTGII PDHPSFSDVG MPPPPAKWKG VCESNFTNKC NNKLIGARSY QLGNGSPIDS
habro GFVSAQKQRM LSLHTTHTPS FLGLQQNMGV WKDSNYGKGV IIGVLDTGIL PDHPSFSDVG MPSPPAKWKG VCESNFTNKC NNKLIGARSY QLGNGSPIDS
corne GFVSAQKQRI LSLHTTHTPS FLGLQQNMGV WKDSNYGKGV IIGVIDTGII PDHPSFSDVG MPPPPAKWKG VCESNFTNNC NNKLIGARSY QLGNGSPIDS
neori GFVSAQKQRI LSLDTTHTPS FLGLQQNMGV WKDSNYGKGV IIGVIDTGII PDHPSFSDVG MPPPPAKWKG VCESNFTNKC NNKLIGARSY QLGNGSPIDS
arcan GFVSAQKQRI LSLDTTHTPS FLGLQQNMGV WKDSNYGKGV IIGVIDTGII PDHPSFSDVG MPPPPAKWKG VCESNFTNKC NNKLIGARSY QLGNGSPIDS
huayl GFVSAQKQRI LSLHTTHTPS FLGLQQNMGV WKDSNYGKGV IIGVIDTGII PDHPSFSDVG MPPPPAKWKG VCESNFTNKC NNKLIGARSY QLGNGSPIDS
chile GFVSAQKQRI LSLHTTHTPS FLGLQQNMGV WKDSNYGKGV IIGVLDTGII PDHPSFSDVG MPPPPAKWKG VCESNFTNKC NNKLIGARSY QLGNGSPIDS
penne GFVSAQKQRI LSLHTTHTPS FLGLQQNMGV WKDSNYGKGV IIGVIDTGII PDHPSFSDVG MPPPPAKWKG VCESNFTNNC NNKLIGARSY QLGNGSPIDS
peruv GFVSAQKQRI LSLHTTHTPS FLGLQQNMGV WKDSNYGKGV IIGVLDTGIL PDHPSFSDVG MPSPPAKWKG VCESNFTNKC NNKLIGARSY QLGNGSPIDS

      201                                                                                                300
lycop IGHGTHTAST AAGAFVKGAN VYGNADGTAV GVAPLAHIAI YKVCNSVGCS ESDVLAAMDS AIDDGVDILS MSLSGGPIPF HRDNIAIGAY SATERGILVS
pimpi IGHGTHTAST AAGAFVKGAN VYGNADGTAV GVAPLAHIAI YKVCNSVGCS ESDVLAAMDS AIDDGVDILS MSLSGGPIPF HRDNIAIGAY SATERGILVS
chees IGHGTHTAST AAGAFVKGAN VYGNADGTAV GVAPLAHIAI YKVCNSVGCS ESDVLAAMDS AIDDGVDILS MSLSGGPIPF HRDNIAIGAY SATERGILVS
habro NGHGTHTAST AAGAFVKGAN VYGNADGTAV GVAPLAHIAI YKVCNSVGCS ESDILAAMDS AIDDGVDILS MSLGGGPIPF HKDNIAIGAY SATERGILVS
corne NGHGTHTAST AAGAFVKGAN VYGNANGTAV GVAPLAHIAI YKVCNSVGCS ESDVLAAMDS AIDDGVDILS MSLGGGPIPF HRDNIAIGAY SATERGILVS
neori NGHGTHTAST AAGAFVKGAN VYGNADGTAV GVAPLAHIAI YKVCNSVGCS ESDVLAAMDS AIDDGVDILS MSLGGGPIPF HRDNIAIGAY SATERGILVS
arcan NGHGTHTAST AAGAFVKGAN VYGNADGTAV GVAPLAHIAI YKVCNSVGCS ESDVLAAMDS AIDDGVDILS MSLGGGPIPF HRDNIAIGAY SATERGILVS
huayl NGHGTHTAST AAGAFVKGAN VYGNANGTAV GVAPLAHIAI YKVCNSVGCS ESDVLAAMDS AIDDGVDILS MSLGGGPIPF HRDNIAIGAY SATERGILVS
chile NGHGTHTAST AAGAFVKGAN VYGNANGTAV GVAPLSHIAI YKVCNSVGCS ESDVLAAMDS AIDDGVDILS MSLGGVPIPF HRDNIAIGAY SATERGILVS
penne NGHGTHTAST AAGAFVKGAN VYGNADGTAV GVAPLAHIAI YKVCNSVGCS ESDVLAAMDS AIDDGVDILS MSLGGGPIPF HRDNIAIGAY SATERGILVS
peruv NGHGTHTAST AAGAFVKGAN VYGNADGTAV GVAPLAHIAI YKVCNSVGCS ESDILAAMDS AIDDGVDILS MSLGGGPIPF HRDNIAIGAY SATERGILVS

      301                                                                                                400
lycop CSAGNSGPSF ITAVNTAPWI LTVGASTLDR KIKATVKLGN GEEFEGESAY RPKISNATFF TLFDAAKNAK DPSETPYCRR GSLTDPAIRG KIVLCSALGH
pimpi CSAGNSGPSF ITAVNTAPWI LTVGASTLDR KIKATVKLGN GEEFEGESAY RPKISNATFF TLFDAAKNAK DPSETPYCRR GSLTDPAIRG KIVLCSALGH
chees CSAGNSGPSF ITAVNTAPWI LTVGASTLDR KIKATVKLGN GEEFEGESAY RPKISNATFF TLFDAAKNAK DPSETPYCRR GSLTDPAIRG KIVLCSALGH
habro CSAGNSGPSL ITAGNTAPWI LTVGASTLDR KIKATVKLGN GEEFEGESAY RPKISNSSFF TLYDAAKNAK DPSKTPYCRR GSLTDPAIRG KIVLCLALGH
corne CSAGNSGPSL VTAGNTAPWI LTVGASTLDR KIKATVKLGN GEEFEGESAY RPKISNATFF TLYDAAKNAK EQSKTPYCKP GSLTDPAIKG KIVLCLAFGD
neori CSAGNSGPSF ITAGNTAPWI LTVGASTLDR KIKATVKLGN GEEFEGESAY RPKISNSTFF TLFDAAKNAK DPSETPYCRR GSLTDPAIRG KIVLCSAFGG
arcan CSAGNSGPSF ITAGNTAPWI LTVGASTLDR KIKATVKLGN GEEFEGESAY RPKISNSTFF TLFDAAKNAK DPSETPYCRR GSLTDPAIRG KIVLCSAFGG
huayl CSAGNSGPSF ITAGNTAPWI LTVGASTLDR KIKATVKLGN GEEFEGESAY RPKISNSTFF TLFDAAKNAK DPSETPYCRR GSLTDPAIRG KIVLCSAFGR
chile CSAGNSGPSL VTAGNTAPWI LTVGASTLDR KIKATVKLGN GEEFEGESAY RPKISNATFF TLFDAAKNAK DPSETPYCRR GSLTDPAIRG KIVLCLAFGG
penne CSAGNSGPSL ITAGNTAPWI LTVGASTLDR KIKSTVKLGN GEEFEGESAY RPKISNSTFF TLFDAAKNAK DPSETPYCKP GSLTDPAIKG KIVLCLAFGG
peruv CSAGNSGPSL ITAGNTAPWI LTVGASTLDR KIRATVKLGN GEEFEGESAY RQKISNSTFF TLFDAAKNAK DPSETPYCRP GSLTDPAIRG KIVLCLAFGR

      401                                                                                                500
lycop VANVDKGQAV KDAGGVGMII INPSQYGVTK SADAHVLPAL VVSAADGTKI LAYMNSTSSP VATIAFQGTI IGDKNAPMVA AFSSRGPSRA SPGILKPDII
pimpi VANVDKGQAV KDAGGVGMII INPSQYGVTK SADAHVLPAL VVSAADGTKI LAYMNSTSSP VATIAFQGTI IGDKNAPMVA AFSSRGPSRA SPGILKPDII
chees VAYVDKGQAV KDAGGVGMII INPSQYGVTK SADAHVLPAL VVSAADGTKI LAYMNSTSSP VATIAFQGTI IGDKNAPMVA AFSSRGPSRA SPGILKPDII
habro VANVDKGQAV KDAGGVGMIV INPSQYGVTK SADAHVLPAL DVSAADGTKI LAYMNSTSNP VATIAFQGTI IGDKNAPMVA AFSSRGPSRA SPGILKPDII
corne VANVDKGQAV KDAGGVGMII INPSQYGVTK SADAHVLPAL VVSAADGTKI LAYMNSTSSP VATIAFQGTI IGDKNAPMVA AFSSRGPSRV SPGILKPDII
neori VTNVDKGQAV KDAGGVGMII INPSQYGVTK SADAHVLPAL VVSAADGTKI LAYMNSTSSP VATIAFQGTI IGDKNAPMVA AFSSRGPSRA SPGILKPDII
arcan VTNVDKGQAV KDAGGVGMII INPSQYGVTK SADAHVLPAL VVSAADGTKI LAYMNSTSSP VATIAFQGTI IGDKNAPMVA AFSSRGPSRA SPGILKPDII
huayl VANVDKGQAV KDAGGVGMII INPSQYGVTK SADAHVLPAL VVSAADGTKI LAYMNSTSSP VATIAFQGTI IGDKNAPMVA AFSSRGPSRA SPGILKPDII
chile VANVDKGQAV KDAGGVGMII INPSQYGVTK SADAHVLPAL VVSAADGTKI LAYMNSTSSP VATIAFQGTI IGDKNAPMVA AFSSRGPSRA SPGILKPDII
penne VANVDKGQAV KDAGGVGMII INPSQYGVTK SADAHVLPAL VVSAADGTKI LAYMNSTSSP VATIAFQGTI IGDKNAPMVA AFSSRGPSRA SPGILKPDII
peruv VANVDKGQAV KDAGGVGMII INPSQYGVTK SADAHVLPAL VVSAADGTKI LAYMNSTSSP VATIAFQGTI IGDKNAPMVA AFSSRGPSRV SPGILKPDII

      501                                                                                                600
lycop GPGANILAAW PTSVDDNKNT KSTFNIISGT SMSCPHLSGV AALLKCTHPD WSPAVIKSAM MTTADTLNLA NSPILDERLL PADIYAIGAG HVNPSRANDP
pimpi GPGANILAAW PTSVDDNKNT KSTFNIISGT SMSCPHLSGV AALLKCTHPD WSPAVIKSAM MTTADTLNLA NSPILDERLL PADIYAIGAG HVNPSRANDP
chees GPGANILAAW PTSVDDNKNT KSTFNIISGT SMSCPHLSGV AALLKCTHPD WSPAVIKSAM MTTADTLNLA NSPILDERLL PADIYAIGAG HVNPSRANDP
habro GPGANILAAW PTSVDDNKNT KSTFNIISGT SMSCPHLSGV AALLKCTHPD WSPAAIKSAM MTTADTLNLA NSPILDERLL PADIYAIGAG HVNPSRANDP
corne GPGANILAAW PTSVDDNKNT KSTFNIMYGT SMSCPHLSGV AALLKCTHPD WSPAAIKSAM MTTADTLNLA NSPILDERLL PADIYAIGAG HVNPSRANDP
neori GPGANILAAW PTSVDDNKNI KSTFNIMSGT SMSCPHLSGV AALLKCTHPD WSPAVIKSAM MTTADTLNLA NSPILDERLL PADIYAIGAG HVNPSRANDP
arcan GPGSNILAAW PTSVDDNKNT KSTFNIISGT SMSCPHLSGV AALLKCTHPD WSPAVIKSAM MTTADTLNLA NSPILDERLL PADIYAIGAG HVNPSRANDP
huayl GPGANILAAW PTSVDDNKNT KSTFNIISGT SMSCPHLSGV AALLKCTHPD WSPAVIKSAM MTTADTLNLA NSPILDERLL PADIYAIGAG HVNPSRANDP
chile GPGANILAAW PTSVDDNKNT KSTFNIMSGT SMSCPHLSGV AALLKCTHPD WSPAVIKSAM MTTADTLNLA NSPILDERLL PADIYAIGAG HVNPSRANDP
penne GSGANILAAW PTSVDDNKNT KSTFNIISGT SMSCPHLSGV AALLKCTHPD WSPAAIKSAM MTTADTLNLA NNPILDERLL PADIYAIGAG HVNPSRANDP
peruv GPGANILAAW PTSVDDNKNT KSTFNIISGT SMSCPHLSGV AALLKYTHPD WSPAAIKSAI MTTADTLNLA NSPILDERLL PADIYAIGAG HVNPSRANDP

      601                                                                                                700
lycop GLVYDTPFED YVPYLCGLKY TDQQVGNLIQ RRVNCSEVKS ILEAQLNYPS FSIFGLGSTP QTYTRTVTNV GDATSSYKVE VASPEGVAIE VEPSELNFSE
pimpi GLVYDTPFED YVPYLCGLKY TDQQVGNLIQ RRVNCSEVKS ILEAQLNYPS FSIFGLGSTP QTYTRTVTNV GDATSSYKVE VASPEGVAIE VEPSELNFSE
chees GLVYDTPFED YVPYLCGLKY TDQQVGNLIQ RRVNCSEVKS ILEAQLNYPS FSIFGLGSTP QTYTRTVTNV GDATSSYKVE VASPEGVAIE VEPSELNFSE
habro GLVYDTPFED YLPYLCGLNY TNRQVGNLLQ RRVNCSEVKS ILEAQLNYPS FSIFGLGSTP QTYTRTVTNV GDATSSYKVE VASPKGVAIE VEPSELNFSE
corne GLVYDTPFED YVPYLCGLKY TDQQVGNLIQ RRVNCSEVKS ILEAQLNYPS FSIFGLGSTP QTYTRTVTNV GDATSSYKVE VASPKGVAVE VEPAELNFSE
neori GLVYDTPFED YVPYLCGLNY TNRQVGNLIQ RRVNCSEVKS ILEAQLNYPS FSIFGLGSTP QTYTRTVTNV GDATSSYKVE VASPKDVVVE VEPAELNFSE
arcan GLVYDTPFED YVPYLCGLNY TNRQVGNLIQ RRVNCSEVKS ILEAQLNYPS FSIFGLGSTP QTYTRTVTNV GDATSSYKVE VASPKEVVVE VEPAELNFSE
huayl GLVYDTPFED YVPYLCGLKY TDQQVGNLIQ RRVNCSEVKS ILEAQLNYPS FSIFGLGSTP QTYTRTVTNV GDATSSYKVE VASPKGVAVE VEPAELNFSE
chile GLVYDTPFED YVPYLCGLKY TDQQVGNLIQ RRVNCSEVKS ILEAQLNYPS FSIFGLGSTP QTYTRTVTNV GDAKSSYKVE VASPKDVVVE VEPAELNFSE
penne GLVYDTPSED YLPYLCGLKY TDQQVGNLLQ RKVNCSEVKS ILEAQLNYPS FSIFGLGSTP QTYTRTVTNV GDATSSYKVE VASPKDVVVE VEPAELNFSE
peruv GLVYDTPFED YVPYLCGLKY TDQQVGNLIQ RRVNCSEVKS ILEAQLNYPS FSIFGLGSTP QTYTRTVTNV GDATSSYKVE VASPKGVAIE VEPSELNFSE

      701                                         745
lycop LNQKLTYQVT FSKTTNSSNP EVIEGFLKWT SNRHSVRSPI AVVSA
pimpi LNQKLTYQVT FSKTTNSSNP EVIEGFLKWT SNRHSVRSPI AVVSA
chees LNQKLTYQVT FSKTTNSSNP EVIEGFLKWT SNRHSVRSPI AVVSA
habro LKQKLTYQVT FSKTSSSSNI EVVEGFLKWT SNRHSVRSPI AVVFA
corne LKQKLTYQVT FSKSSSSSNI EVVEGFLKWT SNRHSARSPI AVVFA
neori LKQKLTYQVT FSKSSSSSNI EVVEGFLKWT SSRHSVRSPI AVVFA
arcan LKQKLTYQVT FSKSSSSSNI EVVEGFLKWT SSRHSVRSPI AVVFA
huayl LKQKLTYQVT FSKSSSSSNI EVVEGFLKWT SNRHSVRSPI AVVFA
chile LKQKLTYQVT FSKSSSSSNI EVVEGFLKWT SNRHSVRSPI AVVLA
penne LKQKLTYQVT FSKTSSSSNI EVVEGFLKWT SNRHSVRSPI AVVFA
peruv LKQKLTYQVT FSKSSSSSNI EVVEGFLKWT SNRHSVRSPI AVVFA
```

**Figure S3** Amino acid alignment of P69B orthologs from wild tomato species.

P69B orthologs were extracted from genomes from wild tomato species and aligned. Residues that are different from P69B of cultivated tomato are highlighted in light grey, dark grey or red for n = 2, 3 and 4 variant residues, respectively. Note that only position 400 has 4 different residues.

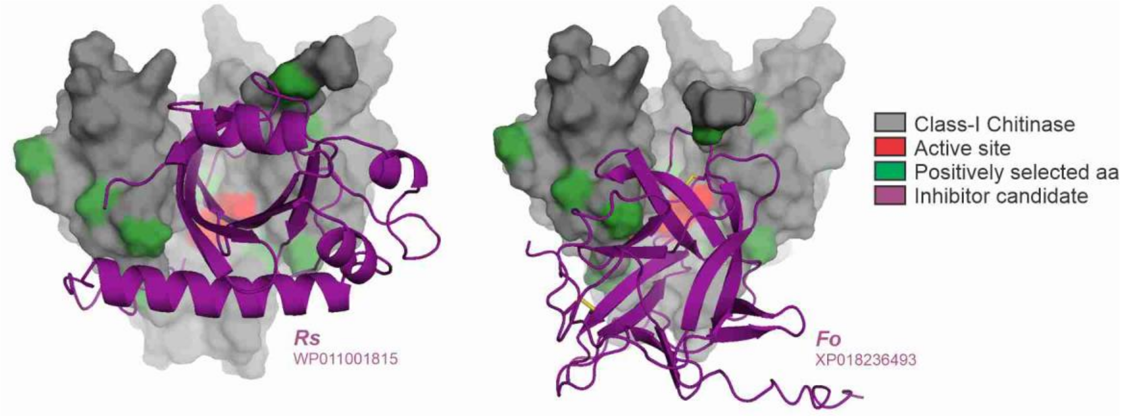


**Figure S4** Two candidate inhibitors interact with positively selected sites in Class-I chitinases. Positively selected sites reported previously (Bishop et al., 2000) were plotted onto the AFM-predicted models of tomato Class-I chitinase in complex with inhibitors from *Ralstonia solanacearum* **(A)** and *Fusarium oxysporum f. sp. lycopersici* **(B)**.

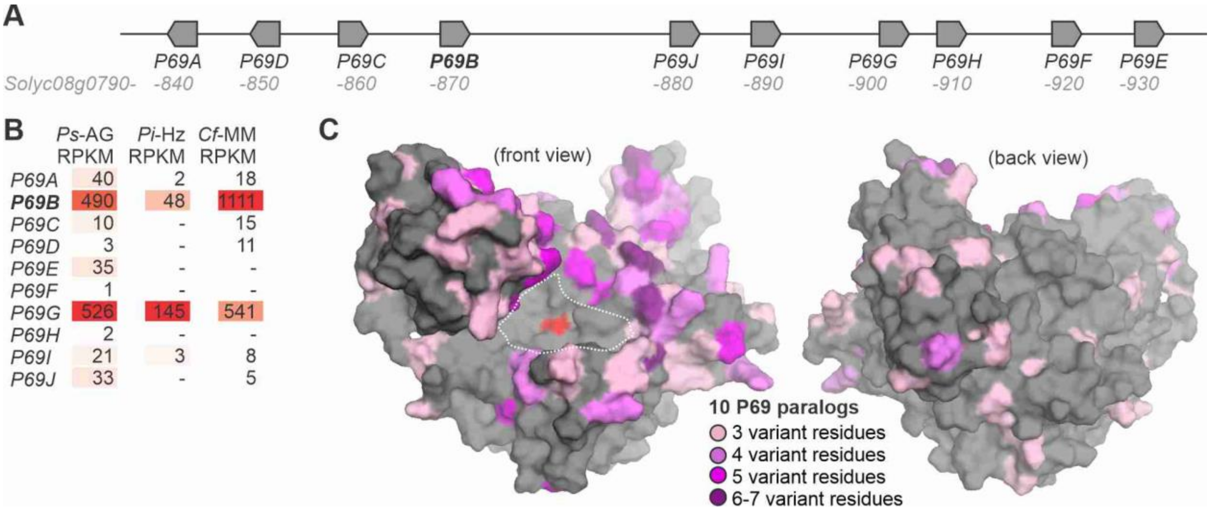


**Figure S5** Expression and variation of the ten P69 paralogs of tomato.

**(A)** Genomic P69 cluster contains ten paralogs. Sequence was extracted from the genome of cv. Heinz (Tomato Genome Consortium, 2012). **(B)** Expression levels in reads per kilobase per million (RPKM) of *P69* genes in three different cultivars infected with *Ps* (cv. Ailsa Graig, Rosli et al., 2013); *Pi* (cv. Heinz, Hwang et al., 2020) and *Cf* (cv. Money Maker, Ilyas et al., 2016). **(C)** Protein sequences of the ten P69 paralogs of tomato (P60A-I) were aligned and the number of variant residues at each position was plotted on the model of the P69B structure. The variation (purple) is surrounding the active site (red) in the substrate binding groove (dashed line) as a 'ring-of-fire'.
