## Supplemental Table for "Alphafold-multimer predicts cross-kingdom interactions at the plant-pathogen interface"

**Supplemental Tables S1-S2-S3**

**Supplemental Table S1** Synthesised nucleotide sequences

**P69B Fragment-1**

GAAGACATAATGGGATTTGTTCTCTTTTCACAATTGCCTTCATTTCTTCTTGTCTCTACACTTCTCTTATTCCTAGTAATATCCCACTCTTGCCGTGCAGGAGGTAATTTAGAGACTTACATCGTCCACGTGGAGAGTCCAGAATCTTTGGTCACCACTCAATCACTCTTGACAGATTTGGGTAGTTACTATCTTTCCTTTCTGCCCAAAACGGCTACTACGATCTCCTCAAGTGGCAACGAGGAAGCCGCGACAATGATTTACTCATATCATAACGTAATGACTGGTTTCGCAGCTAGATTAACTGCTGAGCAGGTCAAAGAAATGGAAAAGAAACATGGATTTGTGTCCGCACAGAAGCAAAGAATCTTGAGCCTACATACTACACACACTCCTAGTTTCCTTGGTTTACAGCAAAATATGGGCGTTTGGAAGGACTCAAATTATGGTAAGGGAGTGATCATTGGTGTAATTGATACAGGTATCATTCCCGACCATCCCAGCTTTAGTGATGTTGGCATGCCCCCGCCACCTGCAAAATGGAAAGGTGTTTGCGAGTCTAATTTTACTAATAAGTGTAATAACAAGCTTATAGGTGCCCGATCTTACCAACTAGGTAATGGTTCCCCAATTGATTCTATAGGGCACGGAACGCACACAGCCAGTACTGCTGCCGGCGCTTTCGTAAAAGGAGCCAACGTGTATGGTAATGCCGACGGAACAGCCGTAGGGGTGGCACCATTAGCTCATATTGCTATTTATAAGGTCTGCAATTCAGTTGGTTGTTCAGAGTCAGACGTACTTGCAGCCATGGACTCCGCAATAGACGATGGTGTTGATATACTTTCTATGTCTTTGTCTGGAGGTCCGATTCCTTTTCACCGGGACAATATCGCTATAGGGGCCTATAGTGCTACTGAACGTGGAATCCTCGTTTCTTGTTCAGCTGGTAATAGTGGACCTTCCTTTATTACAGCTGTCAATACCGCTCCTTGGATTCTAACTGTCGGAGCTTCCACTTTGGACCGTAAGATTAAGGCAACTGTGAAACTGGGCAACGGTGAGGAATTTGAAGGAGAATCAGCATATCGTCCTAAGATAAGCAATGCTACATTCTTTACACTCTTTGATGCGGCAAAGAACGCTAAAGACCCTAGCGAAACCCCCTATTGTCGCAGGGGAAGTCTAACAGACCCTGCTATCAGGGGTAAAATAGTTCTCTGCAGTGCTCTGGGACATGTTGCAAACGTTGATAAAGGACAGGCGGTAAAGGACGCTGGTGGAGTTGGAATGATAATCATAAATCCAAGTCAGTATGGAGTAACCAAGTCTGCTGATGCTCATGTGTTGCCTGCTCTCGTAGTGAGCGCCGCTGATGGTATGTCTTC

**P69B Fragment-2**

GAAGACATTGGTACTAAGATATTGGCATATATGAACTCTACTAGCTCTCCAGTAGCCACAATTGCATTTCAGGGAACGATCATTGGTGACAAGAACGCACCGATGGTTGCGGCATTTAGTTCCCGAGGCCCTTCACGTGCATCCCCTGGTATCCTCAAGCCTGATATCATTGGACCTGGCGCAAACATACTTGCCGCATGGCCTACTTCAGTGGATGACAATAAGAACACAAAGTCTACTTTTAATATAATTAGTGGTACAAGCATGTCCTGTCCCCACCTGTCTGGAGTAGCTGCACTGCTTAAATGTACGCATCCAGATTGGTCACCAGCTGTTATAAAGAGCGCTATGATGACGACCGCTGACACCTTGAACCTCGCAAACTCTCCTATTCTTGACGAGCGTTTGCTTCCAGCTGACATCTATGCAATAGGCGCCGGACATGTGAATCCGAGTAGGGCTAATGACCCAGGGTTGGTGTACGATACTCCTTTCGAAGATTACGTCCCATATCTCTGTGGACTTAAATATACGGATCAACAGGTTGGTAATTTAATTCAAAGACGTGTAAACTGTTCCGAAGTTAAGAGCATCTTGGAAGCTCAACTGAATTATCCCAGTTTTTCTATTTTTGGACTAGGAAGTACTCCTCAGACTTATACAAGAACTGTGACTAATGTCGGAGATGCAACATCTTCCTACAAAGTTGAGGTTGCCAGCCCTGAAGGGGTCGCAATCGAAGTTGAACCTAGCGAGCTTAACTTTTCCGAGCTAAATCAGAAACTTACGTACCAAGTAACATTTTCAAAGACAACTAACTCTAGTAATCCAGAAGTAATCGAGGGTTTTCTTAAATGGACTAGTAATCGTCATAGTGTGCGATCTCCCATCGCCGTTGTAAGCGCAGGTGGGCACCATCACCATCACCACTGAGCTTATGTCTTC

**His-MBP-TEV**

ATGGGCAGCAGCCATCATCATCATCATCATATGAAAATCGAAGAAGGTAAACTGGTAATCTGGATTAACGGCGATAAAGGCTATAACGGTCTCGCTGAAGTCGGTAAGAAATTCGAGAAAGATACCGGAATTAAAGTCACCGTTGAGCATCCGGATAAACTGGAAGAGAAATTCCCACAGGTTGCGGCAACTGGCGATGGCCCTGACATTATCTTCTGGGCACACGACCGCTTTGGTGGCTACGCTCAATCTGGCCTGTTGGCTGAAATCACCCCGGACAAAGCGTTCCAGGACAAGCTGTATCCGTTTACCTGGGATGCCGTACGTTACAACGGCAAGCTGATTGCTTACCCGATCGCTGTTGAAGCGTTATCGCTGATTTATAACAAAGATCTGCTGCCGAACCCGCCAAAAACCTGGGAAGAGATCCCGGCGCTGGATAAAGAACTGAAAGCGAAAGGTAAGAGCGCGCTGATGTTCAACCTGCAAGAACCGTACTTCACCTGGCCGCTGATTGCTGCTGACGGGGGTTATGCGTTCAAGTATGAAAACGGCAAGTACGACATTAAAGACGTGGGCGTGGATAACGCTGGCGCGAAAGCGGGTCTGACCTTCCTGGTTGACCTGATTAAAAACAAACACATGAATGCAGACACCGATTACTCCATCGCAGAAGCTGCCTTTAATAAAGGCGAAACAGCGATGACCATCAACGGCCCGTGGGCATGGTCCAACATCGACACCAGCAAAGTGAATTATGGTGTAACGGTACTGCCGACCTTCAAGGGTCAACCATCCAAACCGTTCGTTGGCGTGCTGAGCGCAGGTATTAACGCCGCCAGTCCGAACAAAGAGCTGGCAAAAGAGTTCCTCGAAAACTATCTGCTGACTGATGAAGGTCTGGAAGCGGTTAATAAAGACAAACCGCTGGGTGCCGTAGCGCTGAAGTCTTACGAGGAAGAGTTGGCGAAAGATCCACGTATTGCCGCCACTATGGAAAACGCCCAGAAAGGTGAAATCATGCCGAACATCCCGCAGATGTCCGCTTTCTGGTATGCCGTGCGTACTGCGGTGATCAACGCCGCCAGCGGTCGTCAGACTGTCGATGAAGCCCTGAAAGACGCGCAGACTAATTCGAGCTCGAACAACAACAACAATAACAATAACAACAACCTCGGGATCGAGGAAAACCTGTATTTTCAGGGCGAATTCGGATCCTCTAGAGTCGACCTGCAGGCAAGCTT

**XpSsp1-WP046932418.1-P3**

GCTCCACCTACAGATACGGCAACTCCCCCCCCAGCGACGGGTTCCACCCGTGGTCCGGCGGCTGGCAAGGCGGTCGTGGTGACCACCACGTGCCGTACCGACGCCGACTGTACCGTGAAGAACGTAGGTAATTGTTGTGGCGCATTTCCGGCGTGCGTGAACGTCAACAGCGCGACCGACCCGAATGGTGTTTTGGCTCAGTGCCAAGCGGGTGGCATGATGAGCGTTTGCGGCTTCCGCGAGATCTCTGCCTGCCAGTGCGTTGCAGGCCAGTGCGCAGCTAAAGATGCCCAAGCGGATACCCTGCGTCCGGCGACTCCGCCGACGGAAACCGTTCACTAG

***Cf*Ecp36-KAH3648627.1-P5**

AAAAGGGGGTCAGGTGGAGAATGTAGAACACAATCCGACTGTTGTTACTGCGACCCGGGTTCCGGCTGCATCCTGAGCTGTCAGGCAGATCGTTCTAGCACCACGGGCTATCGCTGCATTACCAGCCGTCAGTCGGCTTGCAACGTTGCGGGTTGCAGCTGCTAG

***Fo*TIL-XP_018243121.1-P8**

GCTCCCAAGTGTAAAGCAGGAGAACAATTTTTCGAGTGCGGCACCGCTTGCCCGTTGACGTGCGACAAACCGGAACCGCGTCCGTGTACCAAACAATGTGTTCCGGGTTGCTTTTGCAAGGAGGTGCGCATTCTGCGTCAAAGCCAGCAGGTTCTGTCGCCGCTCAAGGCGCTGTTGCTGCCGGTGAGCATCCACAACCGCCCAATCTCTGGTATTTCCCATAGCCGTCACCTGTAG

***Fo*Six15-APP91304.1-P9**

ACAATATATTGTAGGGATGTATCACCACCCCGCGATACCCGTAGCTGGTGTAAAACGAACACCCCGGCGTGGCAAGGTTGCCAGCGTTTTTGCAGCGAGCACTGCCGCTCCACCCCGCGTGACTACCCGGACGGCTGCATGTATCATCTGCAGGTTGGCGGTGATTACGACTGTTTCTGCAAGTAG

**XP001545484.1-P6**

TTCACACCACCCGGATTTGAACCTAGTTCAACGAACGATCTGACCGTCGCATACGGCTCGAAACTTGCCACCAACGGTATTCAAATGCTGCGTGCGGACACCGCGTATGCTCCGATTCTGGGTACAGCGACCAAAATGTCCGGCACCTATGCGGTTATGATGGTTGATCCGGACATCCCGCCGGCAAAGGTGGGTGGTGACACCAGCCAATTTCTGCATTGGATGCAGGCAGACTTGACCAGCTCAAATACTACTACGACCATTGGTGGCCAGAAAATCTACGAGCTGATTAACGTGAAGAATACCTCTGCCTTTGCGACGTACCTGCAGCCGAATCCGCCAGATATCGCTCCGACCACCCACCGTTATACCCAGTTGCTCTTTAACACCACTGGCATGAATATGAGCTTGGCCAGCCTGCAAACGGCGGGTAAAACCCGCGGTAATTTCAACGCGACGAACGTTGTTAAGTCTGCTAAGTTAACGGTGCTGATGGGTAACTCCTTCGACGTGAGCTTCGGCGATAAAGCGATCAACAACACCGGCGCGTCCACCGGCAGCAGCACCTAG

**WP_011000405.1-P7**

GCTGATGATGTTCCCGTAGCGGACGCATCACTCCAACCGAGCGTTGAGGCGGCGCGCCCACCGGTTGCTGCGGCGACCGTCGTGCAAGGTTTTGGCCAGGCAGTTGAGGCAAGCGCACTGGAAGGTTATCGTGGTGGTACGCAGCCGCTGTACCAGACCGTCAACGACGCGCGTTTGTCCGGCACGGTGACCGATAATGCCGCGATTAACGTTGCTACTGGCGCTAATATTGTACGTGACGGCAGCTTCGCCAACGCCTCTGGCATCCCGACCGTTATTCAAAACACCGGTGCGAACGTGCTGATCCAGAATGCGACCATCGTGAACGTGCAGTTCCGCCCGTAG

**Supplemental Table S2** Used plasmids

| **Plasmid** | **Description** | **Reference** |
| --- | --- | --- |
| pJK187 | Binary vector (pL0V2-2x35S::SC) | This work |
| pET-32/28 | Bacterial expression vector | Novinec et al., 2012 |
| P19 | Binary vector carrying p19 silencing inhibitor | Van der Hoorn et al., 2003 |
| pFH20 | Binary vector for P69B-His expression | This work |
| pHJ000 | Bacterial expression vector pET-32/28-His-MBP-TEV | This work |
| pHJ028 | Bacterial expression vector for P3(WP046932418.1) | This work |
| pHJ033 | Bacterial expression vector for P5(KAH3648627.1) | This work |
| pHJ029 | Bacterial expression vector for P6(XP001545484.1) | This work |
| pHJ032 | Bacterial expression vector for P7(WP011000405.1) | This work |
| pHJ030 | Bacterial expression vector for P8(XP_018243121.1) | This work |
| pHJ031 | Bacterial expression vector for P9(APP91304.1) | This work |
| pHJ046 | Bacterial expression vector for Epi1 | This work |
| pHJ047 | Bacterial expression vector for EpiC1 | This work |

**Supplemental Table S3** Primer sequences

| **Primer** | **Use** | **Nucleotide sequence (5’-3’)** |
| --- | --- | --- |
| His-MBP-F | Cloning | AAGAAGGAGATATACCATGGGCAGCAGCCATCATCATCAT  CATCATATG |
| His-MBP-R | Cloning | TGGTGGTGGTGGTGCTCGAGAAGCTTGCCTGCAG |
| Epi1-F | Cloning | CTGTATTTTCAGGGCGAATTCCAAAGCCCGCAAGTCATCAG |
| Epi1-R | Cloning | GACTCTAGAGGATCCGAATTCTTATCCCTCCTGCGGTGTC |
| EpiC1-F | Cloning | CTGTATTTTCAGGGCGAATTCCAAGTGGACGGCGGATACTC |
| EpiC1-R | Cloning | GACTCTAGAGGATCCGAATTCCTACTTAACTGGGGTAATCG |
| P3-F | Cloning | CTGTATTTTCAGGGCGAATTCGCTCCACCTACAGATACGG |
| P3-R | Cloning | GACTCTAGAGGATCCGAATTCCTAGTGAACGGTTTCCGTCGG |
| P5-F | Cloning | CTGTATTTTCAGGGCGAATTCAAAAGGGGGTCAGGTGGAGA |
| P5-R | Cloning | GACTCTAGAGGATCCGAATTCCTAGCAGCTGCAACCCGCAA |
| P6-F | Cloning | CTGTATTTTCAGGGCGAATTCTTCACACCACCCGGATTTGA |
| P6-R | Cloning | GACTCTAGAGGATCCGAATTCCTAGGTGCTGCTGCCGGTGG |
| P7-F | Cloning | CTGTATTTTCAGGGCGAATTCGCTGATGATGTTCCCGTAGC |
| P7-R | Cloning | GACTCTAGAGGATCCGAATTCCTACGGGCGGAACTGCACGT |
| P8-F | Cloning | CTGTATTTTCAGGGCGAATTCGCTCCCAAGTGTAAAGCAGG |
| P8-R | Cloning | GACTCTAGAGGATCCGAATTCCTACAGGTGACGGCTATGGG |
| P9-F | Cloning | CTGTATTTTCAGGGCGAATTCACAATATATTGTAGGGATGT |
| P9-R | Cloning | GACTCTAGAGGATCCGAATTCCTACTTGCAGAAACAGTCGT |
| PJK187-F | Sequencing | CTATCCTTCGCAAGACCCTTC |
| PJK187-R | Sequencing | CTCAACACATGAGCGAAACC |
| pet32a-F | Sequencing | TAATACGACTCACTATAGGG |
| pet32a-R | Sequencing | GCTAGTTATTGCTCAGCGG |
