## Supplemental File S1 for "Alphafold-multimer predicts cross-kingdom interactions at the plant-pathogen interface": README.md

# script-1.py
This script describes the jackhmmer database search as described in AlphaFold2 v2.1.1.

# script-2.py
This script describes calculating mean non-gap MSA depths from AlphaFold multimer output files.

# script-3.py
This script describes generating protein comples `.pse` files for manual inspection.
