## Supplemental File S1 for "Alphafold-multimer predicts cross-kingdom interactions at the plant-pathogen interface": script-1.py

```
#!/usr/bin/env python

## Database search as in AlphaFold v2.1.1, using the reduced dataset and including all databases needed to model multimers

import os
import subprocess
import shutil
import sys
import re


data_dir = sys.argv[1]
fasta_loc = sys.argv[2]
out_dir = sys.argv[3]
n_cpu = sys.argv[4]

jackhmmer_binary = shutil.which("jackhmmer")
n_iter = 1
e_value = 0.0001
filter_f1 = 0.0005
filter_f2 = 0.00005
filter_f3 = 0.0000005

os.system(
    "{} -o /dev/null -A {} --noali --F1 {} --F2 {} --F3 {} --incE {} -E {} --cpu {} -N {} {} {}".format(
        jackhmmer_binary,
        out_dir,
        filter_f1,
        filter_f2,
        filter_f3,
        e_value,
        e_value,
        n_cpu,
        n_iter,
        fasta_loc,
        data_dir,
    )
)
```
