## Supplemental File S1 for "Alphafold-multimer predicts cross-kingdom interactions at the plant-pathogen interface": script-2.py

```
#!/usr/bin/env python

import pickle
import pandas as pd
import numpy as np
import string
from Bio import SeqIO
import glob
import os
import sys
import matplotlib.pyplot as plt
import seaborn as sns
import re
import gzip

# The data_dir can only contain the fasta file used to model the protein complex
data_dir = sys.argv[1]

fasta_loc = glob.glob(os.path.join(data_dir, "*fasta"))[0]
pickle_loc = glob.glob(os.path.join(data_dir, "features.pkl.gz"))[0]

names = []

# extract the names of the protein complex
complex_name = re.search(".*\/([A-Za-z0-9\.]+)_([A-Za-z0-9\.]+)\.fasta", fasta_loc)
names.append(complex_name.group(1))
names.append(complex_name.group(2))

# Read in multi fasta file

print(fasta_loc)
msf = SeqIO.index(fasta_loc, "fasta")
alphabet = string.ascii_uppercase

# Define length of two different chains and their names

chains = {}
for counter, i in enumerate(msf):
    chains[alphabet[counter]] = len(msf[i].seq)


# Open features.pkl file and extract MSA information

with gzip.open(pickle_loc, "rb") as f:
    feat = pickle.load(f)

# Count all gaps (=21) in the MSA

non_gap_coverage = np.count_nonzero(feat["msa"] != 21, axis=0)
x = np.arange(0, len(non_gap_coverage))

# Make df for plotting the different chains

df_gaps = pd.DataFrame(non_gap_coverage, columns=["non_gap_coverage"])
df_gaps.loc[range(0, chains["A"]), "protein"] = names[0]
df_gaps.loc[range(chains["A"], chains["A"] + chains["B"]), "protein"] = names[1]
df_gaps["amino_acid_position"] = df_gaps.index

# Plot non-gap coverage per amino acid position for the different proteins in the complex

plt.figure(figsize=(15, 8))

sns.lineplot(
    data=df_gaps, x="amino_acid_position", y="non_gap_coverage", hue="protein"
).set(
    title="Per-residue count of non-gap amino acids for {}_{}".format(
        names[0], names[1]
    )
)

plt.savefig(os.path.join(data_dir, "msa_coverage_distribution.png"))

# Define different parameters for each chain and the complex and save the file

tot_cov = np.mean(non_gap_coverage)
a_cov = np.mean(non_gap_coverage[: chains["A"]])
b_cov = np.mean(non_gap_coverage[-chains["B"] :])

if a_cov > b_cov:
    ratio = a_cov / b_cov

else:
    ratio = b_cov / a_cov

scores = pd.DataFrame(
    [
        [
            "{}_{}".format(names[0], names[1]),
            names[0],
            names[1],
            tot_cov,
            a_cov,
            b_cov,
            ratio,
        ]
    ],
    columns=[
        "complex_name",
        "A",
        "B",
        "mean_non_gap_coverage_complex",
        "mean_non_gap_coverage_A",
        "mean_non_gap_coverage_B",
        "ratio_mean_coverage_low_over_high",
    ],
)

scores.to_csv(os.path.join(data_dir, "scores_overview.tsv"), index=False, sep="\t")
```
