## Supplemental File S1 for "Alphafold-multimer predicts cross-kingdom interactions at the plant-pathogen interface": script-3.py

```
#!/usr/bin/env python

import pymol
from pymol import cmd
import pickle
import json
import pandas as pd
import numpy as np
import string
from Bio import SeqIO
import glob
import re
import sys
from IPython.utils import io

data_dir = sys.argv[1]
# json file with the host protein and specific colouring commands such as
# coluring the active site in red
path_to_json = sys.argv[2]

fasta_loc = glob.glob("{}/*.fasta".format(data_dir))[0]
pickle_loc = glob.glob("{}/result*.pkl".format(data_dir))[0]
pdb_loc = glob.glob("{}/*.pdb".format(data_dir))[0]


def complex_plddt_scores(name, chain_id, plddt_file):
    cmd.select(name, "chain {}".format(chain_id))
    cmd.alter(name, "b=0")
    cmd.do("run /home/felix/bin/data2bfactor.py")
    cmd.do("data2b_res {}, {}".format(name, plddt_file))
    cmd.do("spectrum b, rainbow_rev, {}, minimum=0, maximum=100".format(name))


def pymol_out(chain_id, capture_file):
    with open("{}/{}_{}_pymol.out".format(data_dir, name, chain_id), "w") as f:
        for lines in capture_file.stdout:
            f.write(lines)


name = re.search(".+\/([a-zA-Z0-9_\.]+)\.fasta", fasta_loc).group(1)

# The first-named protein is considered the host-derived protein
host = re.search("([a-zA-Z0-9\.]+)_", name).group(1)

print(name)

# ## Read and process multi fasta file w/ Biopython

alphabet = string.ascii_uppercase

msf = SeqIO.index(fasta_loc, "fasta")

chains = {}

for counter, i in enumerate(msf):
    chains[alphabet[counter]] = len(msf[i].seq)

with open(pickle_loc, "rb") as f:
    best = pickle.load(f)

df = pd.DataFrame(data=best.get("plddt"))
df[0] = [int(x) for x in df[0]]

for counter, i in enumerate(chains):
    prev = i
    if counter == 0:
        tmp = df[0].head(chains[i])
        tmp.index = range(1, len(tmp) + 1)
        tmp.to_csv("{}/plddt_chain_{}.tsv".format(data_dir, i), header=False, sep="\t")

    else:
        tmp = df[0].tail(chains[i])
        tmp.index = range(1, len(tmp) + 1)
        tmp.to_csv("{}/plddt_chain_{}.tsv".format(data_dir, i), header=False, sep="\t")

df = pd.DataFrame(data=best.get("predicted_aligned_error"))
df[0] = [int(x) for x in df[0]]


# ## Pymol visualisation of per residue plDDt

import pymol
from pymol import cmd

pymol.finish_launching(["pymol", "-qc"])


cmd.load(pdb_loc)

with io.capture_output() as captured_A:
    complex_plddt_scores("target", "A", "{}/plddt_chain_A.tsv".format(data_dir))

with io.capture_output() as captured_B:
    complex_plddt_scores("inh", "B", "{}/plddt_chain_B.tsv".format(data_dir))


cmd.save("{}/{}_ranked_0_plddt_coloured.pse".format(data_dir, name))

pymol_out("A", captured_A)
pymol_out("B", captured_B)


### Colouring of protein complexes for manual processing

pymol.finish_launching(["pymol", "-qc"])

cmd.load(pdb_loc)

with io.capture_output() as captured_B:
    complex_plddt_scores("inh", "B", "{}/plddt_chain_B.tsv".format(data_dir))

    cmd.bg_color("white")
    cmd.do("color grey50, chain A")
    cmd.do("show sticks, (cys/ca+cb+sg) and byres (cys/sg and bound_to cys/sg)")
    cmd.do("show sticks, disulfides")
    cmd.do("show lines")
    cmd.do("color atomic, (not elem C)")
    cmd.do("select don, (elem n,o and (neighbor hydro))")
    cmd.do("select acc, (elem o or (elem n and not (neighbor hydro)))")
    cmd.do("dist HBA, (inh and acc),(target and don), 3.2")
    cmd.do("dist HBD, (inh and don),(target and acc), 3.2")
    cmd.do("delete don")
    cmd.do("delete acc")
    cmd.do("hide (hydro)")
    cmd.do("hide labels,HBA")
    cmd.do("hide labels,HBD")
    with open(path_to_json) as json_file:
        data = json.load(json_file)
        if host in data.keys():
            for i in data[host]:
                cmd.do(data[host][i])


cmd.save("{}/{}_ranked_0_complex_screening.pse".format(data_dir, name))
```
