## Supplementary material for "Alphafold-multimer predicts cross-kingdom interactions at the plant-pathogen interface": Suppplemental File S2

**Supplemental File S2** Protein sequences of inhibitors and proteases used for Figure 1

>P69B *Solanum lycopersicum* Solyc08g079870.3.1 CAA76725.1

MGLLKILLVFIFCSFQWPTIQSNLETYIVHVESPESLVTTQSLLTDLGSYYLSFLPKTATTISSSGNEEAATMIYSYHNVMTGFAARLTAEQVKEMEKKHGFVSAQKQRILSLHTTHTPSFLGLQQNMGVWKDSNYGKGVIIGVIDTGIIPDHPSFSDVGMPPPPAKWKGVCESNFTNKCNNKLIGARSYQLGNGSPIDSIGHGTHTASTAAGAFVKGANVYGNADGTAVGVAPLAHIAIYKVCNSVGCSESDVLAAMDSAIDDGVDILSMSLSGGPIPFHRDNIAIGAYSATERGILVSCSAGNSGPSFITAVNTAPWILTVGASTLDRKIKATVKLGNGEEFEGESAYRPKISNATFFTLFDAAKNAKDPSETPYCRRGSLTDPAIRGKIVLCSALGHVANVDKGQAVKDAGGVGMIIINPSQYGVTKSADAHVLPALVVSAADGTKILAYMNSTSSPVATIAFQGTIIGDKNAPMVAAFSSRGPSRASPGILKPDIIGPGANILAAWPTSVDDNKNTKSTFNIISGTSMSCPHLSGVAALLKCTHPDWSPAVIKSAMMTTADTLNLANSPILDERLLPADIYAIGAGHVNPSRANDPGLVYDTPFEDYVPYLCGLKYTDQQVGNLIQRRVNCSEVKSILEAQLNYPSFSIFGLGSTPQTYTRTVTNVGDATSSYKVEVASPEGVAIEVEPSELNFSELNQKLTYQVTFSKTTNSSNPEVIEGFLKWTSNRHSVRSPIAVVSA

>Pip1 *Solanum lycopersicum* Solyc02g077040.4.1

MASNFFLKNITVVLLLFSILSLYPFIVTSRNLKELSMLERHENWMVHHGRVYKDDIEKEHRFKTFKENVEFIESFNKNGTQRYKLAINKYADLTTEEFTTSFMGLDTSLLSQQESTATTTSFKYDSVTEVPNSMDWRKRGSVTGVKDQGVCGCCWAFSAAAAIEGAYQIANNELISLSEQQLLDCSTQNKGCEGGLMTVAYDFLLQNNGGGITTETNYPYEEAQNVCKTEQPAAVTINGYEVVPSDESSLLKAVVNQPISVGIAANDEFHMYGSGIYDGSCNSRLNHAVTVIGYGTSEEDGTKYWIVKNSWGSDWGEEGYMRIARDVGVDGGHCGIAKVASFPTA

>Epi1 *Phytophthora infestans* PITG_22681

MKSALLFTLVVAAVHAQSPQVISPAPRRESNEIDCPEYCLDVYDPVGDGEGNTYSNECYMKRAKCHNETTPPAWKDLVLITGSSTGEQPPSKKCSTVCPDVELPVCGSNRVRYGNPCELRIAACEHPELNIVEDSGKACVGSKVTPQEG

>EpiC2B *Phytophthora infestans* PITG_09173

MSFLRPTLALLAVTALVTTSAQLNGYSKKEVTPEDTELLQKAQSNVSAYNSDVTSRICYLKVDSLETQVVSGENYKFHVSGCSVNSDKELGGCANQNCESSKYDIVIYSQSWTNTLKVTSITPAN

Highlighted are the predicted signal peptide (grey) and the mature proteins used for AFM modeling (underlined)
