## Supplemental File S4 for "Alphafold-multimer predicts cross-kingdom interactions at the plant-pathogen interface"

**Supplemental File S4** Sequences of six hydrolases used for AFM screen.

> Class I chitinase *Solanum lycopersicum* Solyc09g098540.3.1

MKMRMIMILFLLVIIGGFVHGDENESFRKQPTLVKTVKGTKMCVKDWECNKLSKFCCNLTITDYLDVDQFELLFTKRNSPVAHAVGFWDYGSFIRAAALYQPLGFGTTGGKKMQMKEIAAFFGHVGSKTSCGYGVATGGPLAYGLCYNKEMSPSQDYCDEYFKLTYPCTPGARYYGRGALPIYWNYNYGAIGEALKLNLLDHPEYIEQNATMAFQAAIWRWMNPMKKGQPSAHDAFVGNWKPTKNDTLSKRVPGFGTTMNILYGDGVCGQGDVDAMNNIVSHYLYYLDLMGVGREEAGPHEVLNCAEQKPFNPTAVAAS

> Class II chitinase *Solanum lycopersicum* Solyc05g050130.4.1

VMSIKYGLLYLVLVLLLLSLKLEGGGIVIYWGQNGNEGTLASTCASNNYAIVNIAFLVVFGNGQTPVLNLAGHCNPSVGGCTRLSNDIRTCQSRGIKVMLSLGGGVGSYSLSSADDARNVANYLWNNYLGGQSTSRPLGDAILDGIDFDIERGTTQHWDELARALSGFSQQRKVYLTAAPQCPFPDSWLNGALSTGLFDYVWVQFYNNPPCQYSAGSAVNLKSYWNKWTAIQAGKIFLGLPAAPGAAGSGFIPSDVLVSQVLPAINGSPKYGGVMLWSKFYDNGYSSAIKPRV

> Class V chitinase *Solanum lycopersicum* Solyc07g005090.4.1

STKMANILSCCIVFFLALCSGVMANPHCKGVKGAYYPSWAFSTFPPSSIDTSLFTHIYYAFLVPNNTTFKFDIDDETSKLLFNFTSTLRSSVKTLFSVGGGGEGPARFSRMASTSVSRLSFIKSSIEVARKYKFDGFDLDWEFPQNKKDMENFAILLNEWRVEVKKESLATKRPQLLITAAVYFSVDFFLWGEFRSYPVPSINKNLDWINLMFYDYRGSWDTSATGAQAALFDTKSNVSTSYGLSTWIKAGALRSKLIMGLPLYGRTWKLKDPNVDGIGAPAVGVGPGDEGTLTYREIEKFNEENNAKVVYDSATVSAYSVAGTSWIGFDDTNSVAMKLHYAQSQRLRGYFFWAVAGDKDWKISTTAKQSWIIS

>P69B *Solanum lycopersicum* Solyc08g079870.3.1

MGLLKILLVFIFCSFQWPTIQSNLETYIVHVESPESLVTTQSLLTDLGSYYLSFLPKTATTISSSGNEEAATMIYSYHNVMTGFAARLTAEQVKEMEKKHGFVSAQKQRILSLHTTHTPSFLGLQQNMGVWKDSNYGKGVIIGVIDTGIIPDHPSFSDVGMPPPPAKWKGVCESNFTNKCNNKLIGARSYQLGNGSPIDSIGHGTHTASTAAGAFVKGANVYGNADGTAVGVAPLAHIAIYKVCNSVGCSESDVLAAMDSAIDDGVDILSMSLSGGPIPFHRDNIAIGAYSATERGILVSCSAGNSGPSFITAVNTAPWILTVGASTLDRKIKATVKLGNGEEFEGESAYRPKISNATFFTLFDAAKNAKDPSETPYCRRGSLTDPAIRGKIVLCSALGHVANVDKGQAVKDAGGVGMIIINPSQYGVTKSADAHVLPALVVSAADGTKILAYMNSTSSPVATIAFQGTIIGDKNAPMVAAFSSRGPSRASPGILKPDIIGPGANILAAWPTSVDDNKNTKSTFNIISGTSMSCPHLSGVAALLKCTHPDWSPAVIKSAMMTTADTLNLANSPILDERLLPADIYAIGAGHVNPSRANDPGLVYDTPFEDYVPYLCGLKYTDQQVGNLIQRRVNCSEVKSILEAQLNYPSFSIFGLGSTPQTYTRTVTNVGDATSSYKVEVASPEGVAIEVEPSELNFSELNQKLTYQVTFSKTTNSSNPEVIEGFLKWTSNRHSVRSPIAVVSA

>Pip1 *Solanum lycopersicum* Solyc02g077040.4.1

MASNFFLKNITVVLLLFSILSLYPFIVTSRNLKELSMLERHENWMVHHGRVYKDDIEKEHRFKTFKENVEFIESFNKNGTQRYKLAINKYADLTTEEFTTSFMGLDTSLLSQQESTATTTSFKYDSVTEVPNSMDWRKRGSVTGVKDQGVCGCCWAFSAAAAIEGAYQIANNELISLSEQQLLDCSTQNKGCEGGLMTVAYDFLLQNNGGGITTETNYPYEEAQNVCKTEQPAAVTINGYEVVPSDESSLLKAVVNQPISVGIAANDEFHMYGSGIYDGSCNSRLNHAVTVIGYGTSEEDGTKYWIVKNSWGSDWGEEGYMRIARDVGVDGGHCGIAKVASFPTA

> AED1 *Solanum lycopersicum* Solyc08g067100.2.1

MKGLFVKYPNTIIVSPMATRSVLFSSSFLLILLSFSLEKSNALEGRKTIESNFHTIQLTSILPSSSCKPSSKGKRGGASLEVINKHGPCSQLNKKGEKGPTLTEMLAHDQARVDSIQTRIAAQNFNLFRKTEKTSKKYRAKDSKTTLPAQPGIALSTGNYIVTVGIGTPKKDLTLIFDTGSDLTWTQCEPCFKTCFPQQQPIFNPSSSSTYSNISCSSTACSGLKSATGNSPVCSSSTCVYGIQYGDSSFSIGFFAKDRLTLSATDVFDGFMFGCGQDNKGLFGKTAGLIGLGRDPLSIVSQTSAKFGKYFSYCLPTRRGSNGHLSFGKNGAKSNLQFTPFASSQGTSFYFIDVLGISVGGKSLAISPMVFKNAGTIIDSGTVITRLPSTAYSNLRATFREFMSKYPRAPDLSLLDTCYDLSNYTTISIPKISFNFNGNTKMDIVPNGIFIVNGASQVCLAFAGNGDDDSIGIFGNTQQQTMEIVYDVAGEKLGFGYGGCT

Highlighted are the predicted signal peptide (grey) and the mature proteins used for AFM modeling (underlined)
